# Dynamin-2 Phosphorylation as A Critical Regulatory Target of Bin1 and GSK3α for Endocytosis in Muscle

**DOI:** 10.1101/2021.10.11.463889

**Authors:** Jessica Laiman, Julie Loh, Wei-Chun Tang, Mei-Chun Chuang, Hui-Kang Liu, Bi-Chang Chen, Yi-Cheng Chang, Lee-Ming Chuang, Ya-Wen Liu

## Abstract

Tight regulation of endocytosis ensures accurate control of cellular signaling and membrane dynamics, which are crucial for tissue morphogenesis and functions. Mutations of Bin1 and dynamin-2 (Dyn2), proteins that generate membrane curvature and sever endocytic invaginations, respectively, cause progressive hereditary myopathy. Here, we show that Bin1 inhibits Dyn2 via direct interaction of its SRC Homology 3 (SH3) domain with the proline-rich domain (PRD) of Dyn2. Phosphorylation of S848 of Dyn2 by GSK3*α*, a kinase downstream of insulin signaling, relieves Dyn2 from the inhibition of Bin1 and promotes endocytosis in muscle. Mutations of Bin1 associated with centronuclear myopathy disrupt its inhibition of Dyn2, thereby exaggerating Dyn2 fission activity and causing excessive fragmentation of T-tubules in the muscle cells. Our work reveals how Bin1-Dyn2 interaction fine-tunes membrane remodeling at the molecular level, and lay the foundation for future exploration of endocytic regulation and hereditary muscle diseases.

## INTRODUCTION

Endocytosis controls substance uptake, membrane dynamics and signal transduction, and is critical for the structure and function of eukaryotic cells (Conner and Schmid, 2003; Kumari et al., 2010; McMahon and Boucrot, 2011). The importance of endocytosis is exemplified by centronuclear myopathy (CNM), a hereditary muscle disease characterized by centered nuclei, disorganized plasma membrane invaginations (T-tubules) and progressive atrophy of skeletal muscles, and its association with mutations of genes encoding endocytic machineries (Cowling et al., 2012; Dowling et al., 2008; Koch et al., 2021). CNM mutations are found in *DNM2* and *BIN1*, which encode Dynamin 2 (Dyn2) and Bridging integrator 1 (Bin1/amphiphysin 2), two crucial proteins that mediate endocytosis (Bitoun et al., 2005; Nicot et al., 2007). A thorough understanding of the molecular mechanisms through which these endocytic proteins interact and regulate membrane remodeling thus offers an excellent opportunity to elucidate the pathogenesis of CNM and endocytic regulation in muscle.

Dyn2 is a large GTPase that mediates vesicle release from the plasma membrane in many forms of endocytosis (Ferguson and De Camilli, 2012; McMahon and Boucrot, 2011; Mettlen et al., 2009). Through the interaction of its C-terminal proline-arginine rich domain (PRD) with SRC Homology 3 (SH3) domain-containing proteins, Dyn2 is recruited to the plasma membrane, where it binds directly to phosphatidylinositol-4,5-bisphosphate (PI(4,5)P_2_) via its pleckstrin homology (PH) domain (Fig. 1 A) (Grabs et al., 1997; Meinecke et al., 2013; Zheng et al., 1996), induces conformational change that facilitates its self-assembly, GTP hydrolysis and membrane fission activity (Chappie et al., 2010; Faelber et al., 2011; Kong et al., 2018). CNM-associated mutations of Dyn2 increase its activity and result in the fragmentation of T-tubules in both insects and vertebrates (Chin et al., 2015; Cowling et al., 2011; Gibbs et al., 2014; Kenniston and Lemmon, 2010; Wang et al., 2010). This suggests that Dyn2 activity is under tight regulation, through its interacting proteins and auto-inhibition, to ensure the integrity of membrane structure and functions, yet the molecular actions remain incompletely understood (Al-Qusairi and Laporte, 2011; Hohendahl et al., 2016).

**Figure 1.**
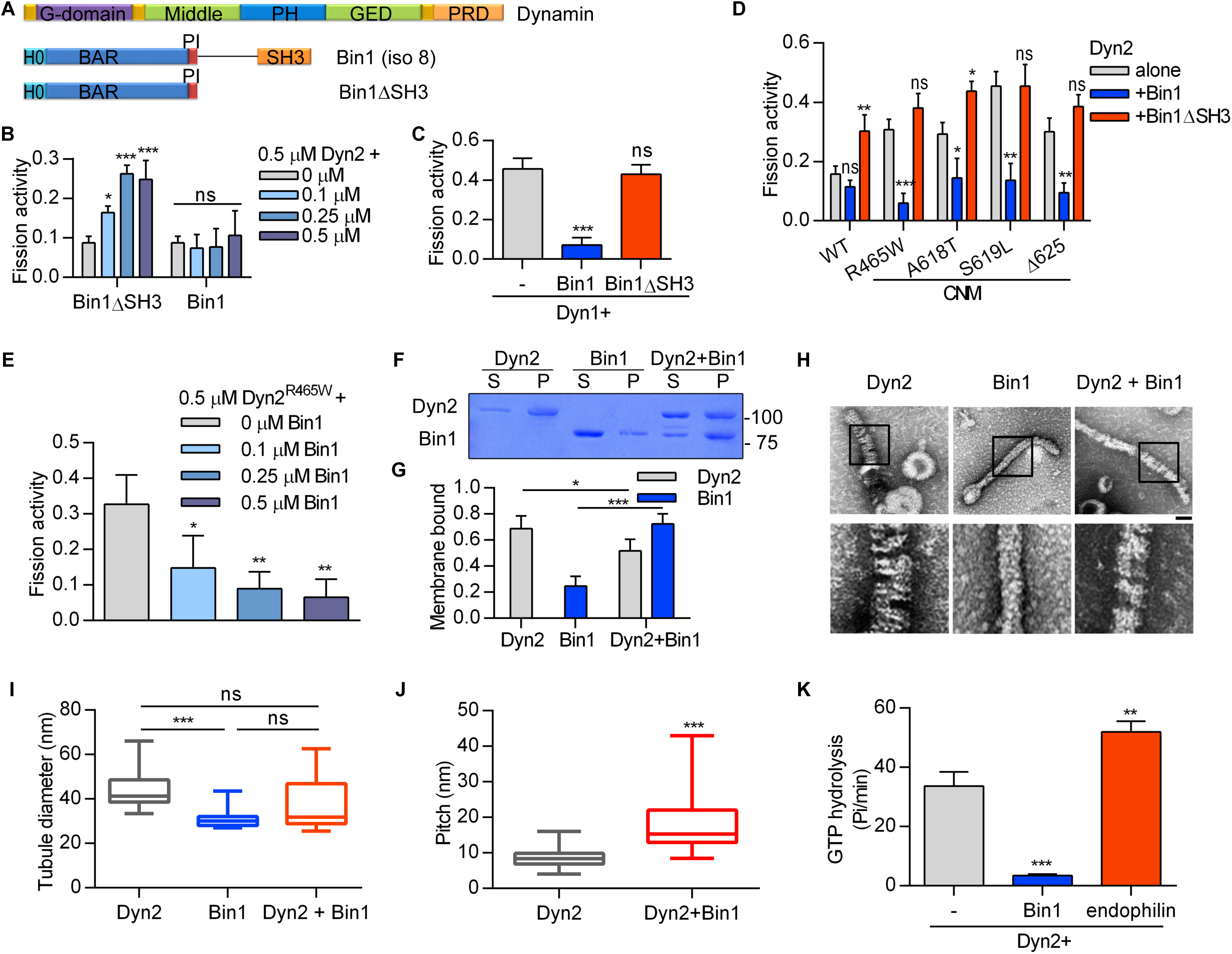
Bin1 inhibits Dyn2 fission activity via SH3 domain. (A) Domain structure of Dynamin and Bin1 constructs used in this study. (B) *In vitro* fission assay of Dyn2. Purified Dyn2 was incubated with increasing concentrations of Bin1 or Bin1*λλ*SH3 in the presence of GTP and SUPER template. The fission activity was measured as the fraction of vesicle released from total SUPER template (n = 3). (C-E) Effect of Bin1 on the fission activity of dynamins. Purified Dyn1 (C) or Dyn2 CNM mutants (D) is incubated with Bin1, Bin1*λλ*SH3 or increasing concentrations of Bin1 (E) (n = 3). (F, G) Liposome binding ability. Dyn2 or Bin1 were incubated with 100-nm liposome for 10 min at 37°C. Liposome-bound proteins (pellet, p) were separated from unbound ones (supernatant, s) with centrifugation sedimentation. The fraction of liposome-bound proteins were quantified and shown in (G) (n = 4). (H-J) Assembly of Dyn2 or Bin1 on liposome. Dyn2 and/or Bin1 were incubated with liposome and then visualized with negative stain TEM. Scale, 100 nm. Boxed areas were magnified and shown as below. The diameter of tubulated membrane were quantified and shown in (I) (n ≥ 11). The periodicity of Dyn2 or Dyn2-Bin1 spirals assembled on liposomes were quantified and shown in (J) (n ≥ 35). (K) Liposome stimulated GTPase activity of Dyn2. Dyn2 was incubated alone or with Bin1 or Endophilin in the presence of liposome and GTP at 37°C. The rate of GTP hydrolysis was measured using a colorimetric malachite green assay (n = 3). Bar graphs are shown as average ± SD, box plots are shown as median ± min and max value. Data are analyzed with one-way ANOVA except for (J) that is analyzed with Student’s t test. ns, no significance; *P< 0.05; **P< 0.01; ***P< 0.001.

Bin1 contains N-BAR and SH3 domains that generates membrane curvature and binds the PRD of dynamins, respectively (Fig. 1 A) (Owen et al., 1998; Peter et al., 2004). Bin1 is essential for the biogenesis and maintenance of T-tubules in skeletal muscle of *Drosophila*, mouse and human, and mutations of Bin1 are associated with CNM (Al-Qusairi and Laporte, 2011; Böhm et al., 2013; Lee et al., 2002; Razzaq et al., 2001). Three observations suggest that Bin1 is a negative regulator of Dyn2. First, Dyn2 severs and Bin1 extends the membrane (Chin et al., 2015; Lee et al., 2002). Second, CNM-associated Dyn2 mutations are hypermorphic, whereas CNM-associated Bin1 mutations reduce its function (Chin et al., 2015; Kenniston and Lemmon, 2010; Wu et al., 2014). Third, downregulation of Dyn2 rescues the phenotypes of the Bin1-related CNM mouse model (Cowling et al., 2017; Rabai et al., 2019). Thus, we aim to decipher the inhibitory and relieving mechanisms of Bin1-Dyn2 interaction.

Here we show that Bin1 inhibits Dyn2 activity through its SH3 domain, which CNM mutations are found to alter. We map a critical serine residue, S848, of Dyn2 that governs the physical interaction between Bin1-SH3 and Dyn2-PRD. Our results suggest that phosphorylation of Dyn2-S848 by glycogen synthase kinase 3α (GSK3α), a component of the insulin signaling pathway, relieves Dyn2 from the inhibition by Bin1. We demonstrate the implications of this Bin1-Dyn2 interplay for tight regulation of endocytosis and pathogenesis of CNM.

## RESULTS

### Bin1 inhibits the fission activity of Dyn2

To investigate the effect of Bin1 on Dyn2 fission activity, we utilized an *in vitro* membrane fission assay to measure Dyn2-severed vesicles released from the supported bilayer with excess membrane reservoir (SUPER) templates (Pucadyil and Schmid, 2008) (Fig. 1 B). As previously shown for the N-BAR domain of endophilin (Neumann and Schmid, 2013), the N-BAR domain along with the PI motif of Bin1 (Bin1ΔSH3) enhanced Dyn2-mediated membrane fission in a concentration-dependent manner. By contrast, full-length Bin1 had no effect (Fig. 1 B). To test whether Bin1 inhibits the fission activity of dynamins, we checked its effects on membrane fission activity of Dyn1 and CNM mutants of Dyn2, both of which possess higher membrane fission activity than WT-Dyn2 as a result of stronger curvature generation and enhanced self-assembly, respectively (Chin et al., 2015; Liu et al., 2011a). As expected, these proteins exhibited greater fission activity than WT-Dyn2 (Fig. 1, C and D), and were inhibited by Bin1 in a dose-dependent manner (Fig. 1 E). These results reveal dual roles of Bin1 on Dyn2 fission activity governed by the SH3 domain: in the absence of the SH3 domain, Bin1 stimulates Dyn2, and the inclusion of SH3 strongly inhibits membrane fission activity of Dyn2.

The inhibitory effect of Bin1 on Dyn2 does not arise from interfering with membrane binding of Dyn2, as revealed by liposome binding assays in which substantial amount of Dyn2 remained bound to the membrane in the presence of Bin1 (Fig. 1, F and G). By contrast, incubation of Bin1 with Dyn2 enhanced membrane binding of Bin1, suggesting that interaction with Dyn2 relieves Bin1 from intramolecular autoinhibition that occurs between the PI motif and SH3 domains (Kojima et al., 2004; Wu et al., 2014). To further understand how Dyn2 and Bin1 assemble on the membrane, we performed negative-stain transmission electron microscope (TEM) to image Dyn2 and Bin1 proteoliposomes. Dyn2 assembled on liposomes of 100 nm diameter resulted in membranous tubule formation with an average diameter of 44.5 ± 2.3 nm and a spiral appearance regularly spaced with pitch of 8.46 ± 2.31 nm (Fig. 1, H-J). Assembly of Bin1 on the liposomes also induced the formation of membrane tubules of a smaller diameter (31.4 ± 1.2 nm) that lack the regular spacing pattern observed with Dyn2. The presence of both Dyn2 and Bin1 led to fewer membrane tubules of an intermediate tubule diameter (37.4 ± 3.7 nm), with less-ordered, spiral-like protein assembly (pitch = 18.35 ± 8.10 nm). This is consistent with a functional antagonism between Dyn2 and Bin1 in membrane remodeling, which is supported by the result that Bin1 also inhibits Dyn2 GTPase activity in the presence of liposomes (Fig. 1 K). By contrast, endophilin, an SH3 domain-containing protein that binds Dyn2, promoted the GTPase activity of Dyn2 in the liposome assays (Fig. 1 K), indicating that distinct SH3 domains have specific regulatory effects on Dyn2. These results lead us to hypothesize that the SH3 domain of Bin1 inhibits Dyn2 fission activity via altering Dyn2 assembly on the membrane.

### PI motif of Bin1 does not affect membrane fission activity of Dyn2

The muscle specific isoform of Bin1 contains a unique positively-charged, 15-amino acid stretch called the PI motif, which enhances the lipid binding ability of Bin1 (Lee et al., 2002). Given that membrane-bound Bin1 inhibits Dyn2 activity, we tested the potential role of PI motif in Dyn2-mediated membrane fission assay using different Bin1 variants (Fig. 2A). To our surprise, PI motif did not affect Bin1 regulation on Dyn2 fission activity, despite the increase of binding affinity between Dyn2 and Bin1 without PI motif (Bin1*λλ*PI) (Fig. 2, B-D) (Kojima et al., 2004; Wu et al., 2014). Considering that Bin1 is important for the formation of T-tubules, which are enriched with cholesterol and sphingomyelin resulting in higher membrane rigidity (Rosemblatt et al., 1981; Roux et al., 2005), we added cholesterol and different percentages of brain sphingomyelin (BSM) to our lipid template. Even on these stiffer membranes, the PI motif did not significantly affect Dyn2 fission activity (Fig. 2 E). Nonetheless, and consistent with a previous report (Lee et al., 2002), we found that the PI motif significantly improved the binding of Bin1 N-BAR domain to membrane (Fig. 2, F and G). In accordance with its ability to promote membrane binding, the PI motif gave rise to greater membrane tubulation of SUPER templates *in vitro* (Fig. 2, H and I) and of plasma membrane in cells (Fig. 2 J). These results suggest that the PI motif is dispensable for regulating Dyn2 fission activity but important for membrane binding and tubulation of Bin1.

**Figure 2.**
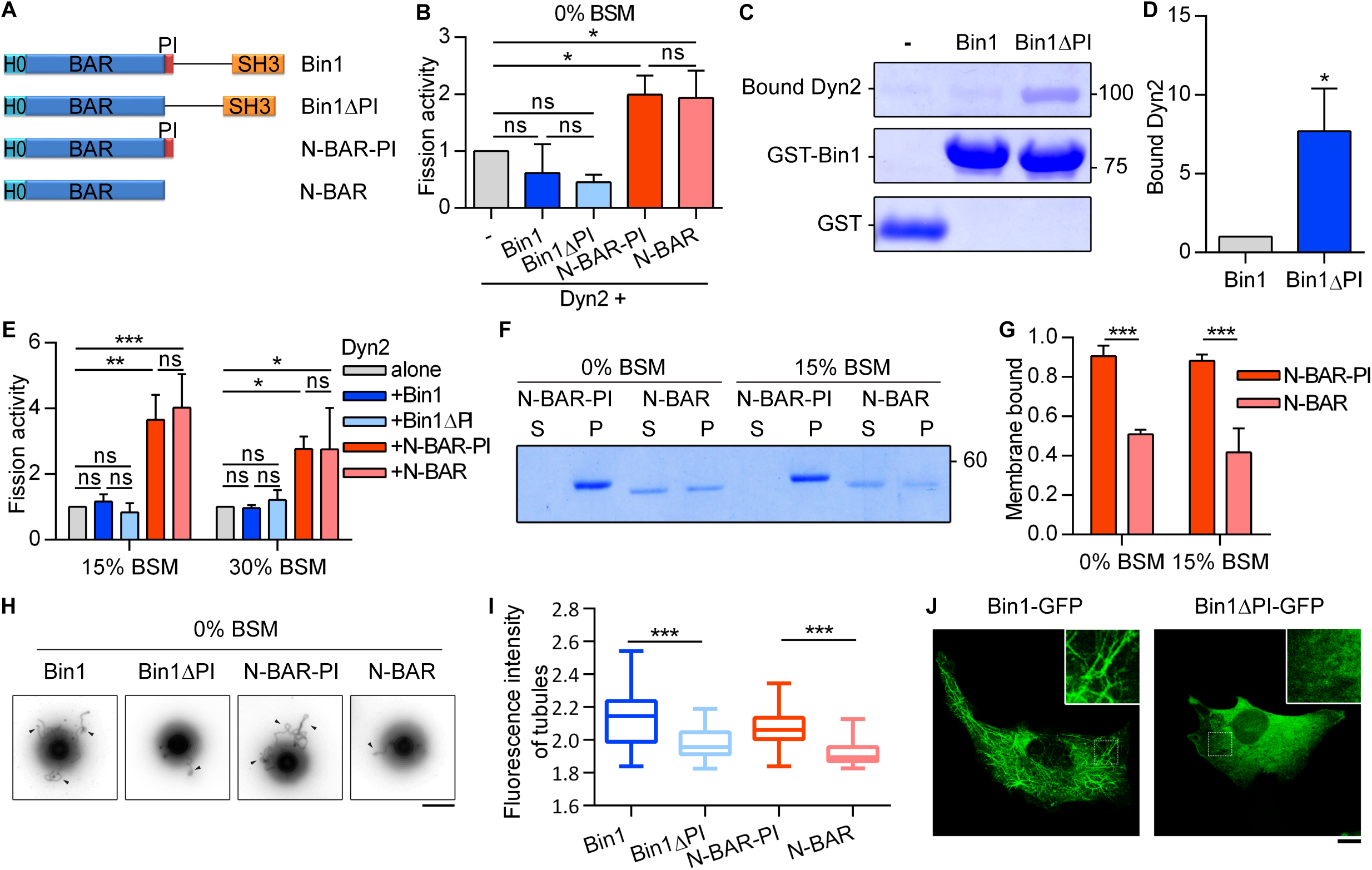
PI motif is important for membrane binding and tubulation of Bin1. (A) Domain structure of Bin1 variants used in this study. (B) Effect of PI motif from Bin1 on regulating Dyn2 fission activity. 0.5 µM purified Dyn2 was incubated by itself or together with 0.5µM Bin1 variants with or without the PI motif in the presence of GTP and SUPER template with lower membrane rigidity. The data was shown as fold change relatively to the Dyn2 fission activity alone (n = 3). (C, D) Effect of PI motif on Bin1-Dyn2 interaction. Different GST-tagged Bin1 proteins (0.5 µM) were incubated with 0.3 µM Dyn2, then the bound Dyn2 was detected and quantified by SDS-PAGE electrophoresis and CBR staining. The data was quantified and shown in (D) (n = 3). (E) Effect of PI motif from Bin1 on regulating Dyn2 fission activity on membrane with higher rigidity. 0.5 µM purified Dyn2 was incubated by itself or together with 0.5 µM Bin1 variants with or without the PI motif in the presence of GTP and SUPER template supplied with 50% cholesterol and indicated percentage of Brain Sphingomyelin (BSM) (n = 3). (F) Effect of PI motif on Bin1 membrane binding ability. N-BAR domain of Bin1 with PI motif (N-BAR-PI) or N-BAR domain are incubated with 100 nm liposome containing indicated percentage of BSM. Liposome-bound proteins (pellet, p) were separated from unbound ones (supernatant, s) with centrifugation sedimentation. The ratio of liposome-bound proteins were quantified and shown in (G) (n = 4). (H) Effect of PI motif on Bin1 membrane tubulation ability *in vitro*. SUPER template was added with 0.5 µM Bin1 variants for 10 min at room temperature and viewed under fluorescence microscope. Tubules from SUPER template were marked by black arrowheads. Scale, 5 µm. The average fluorescence intensity of SUPER template tubules generated by each Bin1 variants was quantified and shown in (I) (n ≥ 36). (J) Effect of PI motif on membrane tubulation ability of Bin1 *in cellulo*. GFP-tagged Bin1 containing PI motif (Bin1-GFP) or without PI motif (Bin1ΔPI-GFP) were over-expressed in C2C12 myoblasts through transfection and viewed under confocal microscopy. Scale, 10 µm. Bar graphs are shown as average ± SD, box plots are shown as median ± min and max value. Data are analyzed with one-way ANOVA (B, E, G, and I) or Student’s t-test (D). ns, no significance; *P< 0.05; ***P< 0.001.

### CNM-associated Bin1 with SH3 domain mutation enhances Dyn2 fission activity

Among the CNM-related Bin1 mutations, Bin1^Q434X^ and Bin1^K436X^ contain premature stop codons that truncate the SH3 domain (Fig. 3 A) (Bohm et al., 2010; Nicot et al., 2007). We speculate these CNM-Bin1 mutants may lose the inhibitory effect on Dyn2. Indeed, GST pulldown assay showed significant decrease in the binding of Bin1^Q434X^ and Bin1^K436X^ to GST-Dyn2PRD (Fig. 3, B and C), while displaying membrane binding and tubulation activities similar to those of Bin1^WT^ (Fig. S1, A-C). Bin1^K436X^ significantly increased Dyn2 fission activity *in vitro* and enhanced tubule formation by Dyn2 (Fig. 3, D-E and S1 D), while its dissociation constant (K_d_, 267 ± 70 nM) was not very different from that of Bin1^WT^ (180 ± 60 nM) (Fig. S1, E and F). These data support the idea that Bin1-SH3 inhibits Dyn2, and enhanced Dyn2 activity could be part of the pathogenesis of CNM associated with Bin1 mutations.

**Figure 3.**
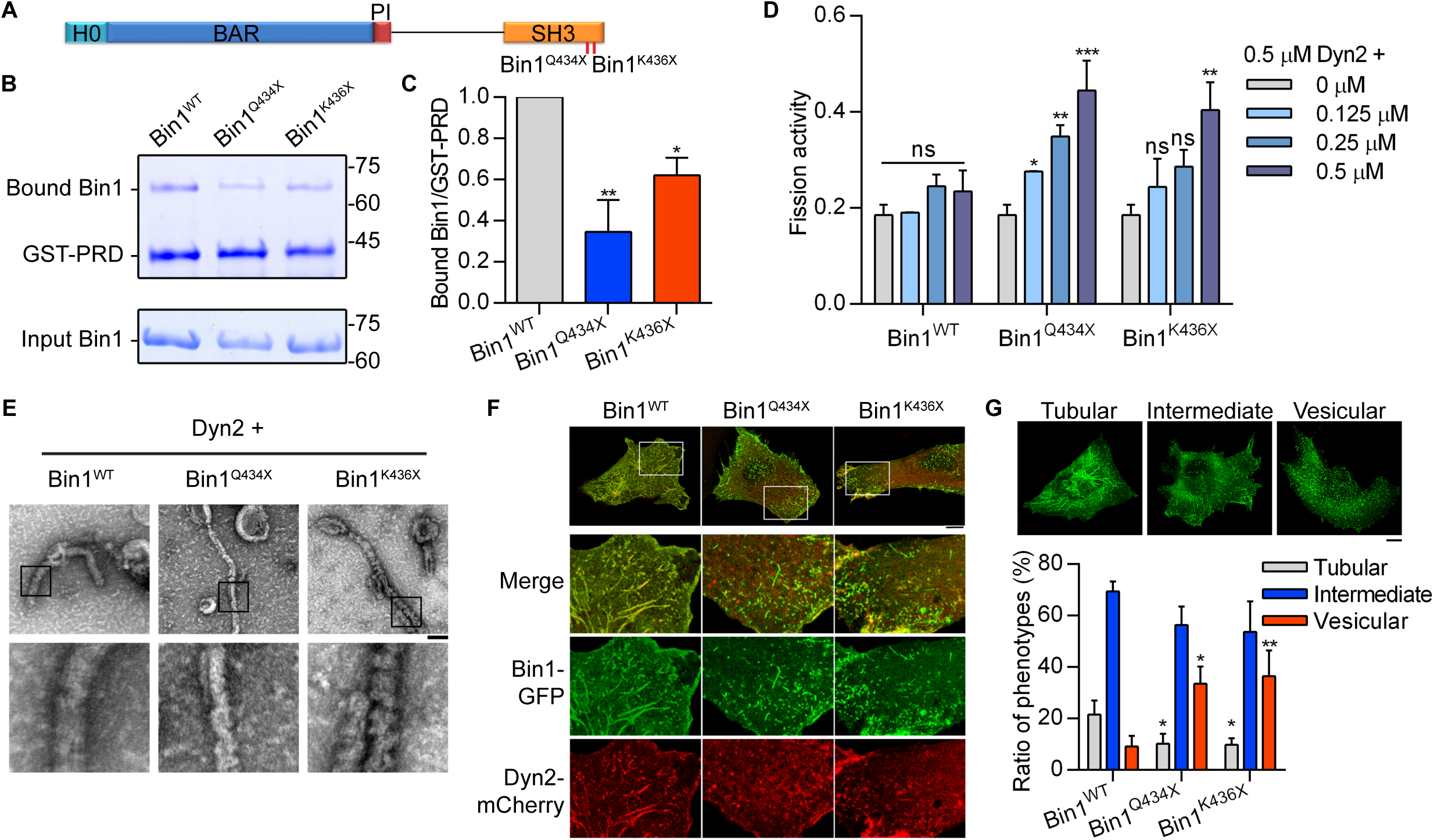
CNM-Bin1 mutants enhance Dyn2 fission activity. (A) CNM-associated Bin1 mutants used in this study. (B-C) Binding ability between Bin1 mutants and the PRD of Dyn2. GST-PRD was incubated with indicated purified His-tagged Bin1 in the presence of liposome. PRD-bound Bin1 was analyzed with SDS-PAGE and Coomassie Blue staining. The ratio of bound Bin1 was normalized to wild-type and shown in (C) (n = 3). (D) Effects of CNM-Bin1 on Dyn2 fission activity. Dyn2 was incubated with increasing concentrations of Bin1^WT^, Bin1^Q434X^, or Bin1^K436X^ in the presence of GTP and SUPER template. The fission activity was determined by sedimentation and fraction of released fluorescent vesicles in the supernatant from total SUPER template (n = 3). (E) Electron micrographs of Dyn2 together with Bin1 mutants assembled onto liposomes. Wild-type Dyn2 and Bin1 mutants were incubated with 100-nm liposomes for 10 min at 37° C, adsorbed to grids and imaged by negative-stain TEM. Scale, 100 nm. Boxed areas were magnified and shown. (F-G) Morphology of CNM-associated Bin1 mutants co-expressed with Dyn2-mCherry in myoblast. GFP-tagged wild-type or mutant Bin1 was transfected into C2C12 myoblasts together with Dyn2-mCherry. The Bin1-mediated membrane morphologies were imaged with confocal microscopy. Bottom panels, magnified images from insets in top panels. The Bin1-mediated membrane deformation was categorized into three groups (the examples of cell from each category are shown in top panel of (G)), and the ratio of each population was quantified and compared with Bin1^WT^ as shown in (G) (n ≥ 75 from three independent repeats). Bar graphs are shown as average ± SD, box plots are shown as median ± min and max value. Data are analyzed with one-way ANOVA. ns, no significance; *P< 0.05; **P< 0.01; ***P< 0.001.

Consistent with the result from *in vitro* membrane tubulation assays (Fig. S1 C), expression of GFP-tagged Bin1^Q434X^ and Bin1^K436X^ in the C2C12 myoblasts induced tubulation of the plasma membrane similar to that by Bin1^WT^-GFP (Fig. S1 G). The morphology of these membrane tubules were reminiscent of T-tubules in terms of their high curvatures and Bin1 enrichment, and they could be severed into small vesicles by Dyn2 (Chin et al., 2015; Laiman and Liu, 2020), making them a good cellular system to validate Bin1-Dyn2 interactions observed in the reconstitution experiments. We transfected both Dyn2-mCherry and Bin1-GFP into the C2C12 myoblasts and examined the morphology of membrane tubules (Fig. 3 F). We categorized the cells based on the overall Bin1-GFP tubulation patterns (Fig. 3, F and G). Bin1^Q434X^ or Bin1^K436X^ significantly reduced the proportion of cells predominated with Bin1 tubules, with a reciprocal increase in cells that showed the characteristic vesicular membrane patterns (Fig. 3 G). This result is consistent with higher Dyn2 fission activity in the presence of Bin1^Q434X^ or Bin1^K436X^ compared to Bin1^WT^. Furthermore, live cell imaging of the dynamics of Bin1-mediated tubules revealed more frequent fission events in the presence of Bin1^Q434X^ or Bin1^K436X^ (Video S1 and Fig. S1 H). Together, these data suggest that CNM-associated Bin1 mutants with truncated SH3 domain have weakened inhibition on Dyn2, underscoring the importance of Bin1-SH3 in tuning Dyn2 fission activity.

### Phosphorylation of Dyn2 PRD confers resistance to Bin1 inhibition and promotes endocytosis

The above data with CNM-associated Bin1 mutations indicate that Dyn2-PRD could be a regulatory hub for Dyn2 activity under physiological contexts. Given that phosphorylation of Dyn1 PRD has been known to regulate Dyn1 function (Clayton et al., 2010; Reis et al., 2015; Tan et al., 2003), we hypothesized that similar mechanisms operate in Dyn2-PRD. Among the putative phosphorylation sites on Dyn2-PRD, we focused on Ser848 and Ser856, two serine residues adjacent to the SH3 binding pocket of Dyn2 PRD (Fig. 4 A) (Choudhary et al., 2009; Efendiev et al., 2002). Using GST pulldown assays, we found that Dyn2^S848E^, a phosphomimetic mutant of Dyn2-S848, had decreased binding affinity to Bin1-SH3, but not to endophilin-SH3 (Fig. 4 B, C and S2 A-C). *In vitro* fission assays and analysis of Bin1-GFP tubule in C2C12 myoblasts further confirmed that the membrane fission activity of Dyn2^S848E^ was increased by Bin1 (Fig. 4 D, E, S2D and Video S2), reminiscent of that of Dyn2-WT with Bin1ΔSH3 (Fig. 1 B). Moreover, Bin1 and Dyn2 ^S848E^ assembled into smaller tubules on liposomes similar to those of Dyn2^WT^ and Bin1^Q434X^ or Bin1^K436X^ (Fig. 4, F and G). These results raise the possibility that Dyn2-S848 is a critical residue that governs Dyn2-Bin1 interaction, likely through its phosphorylation status.

**Figure 4.**
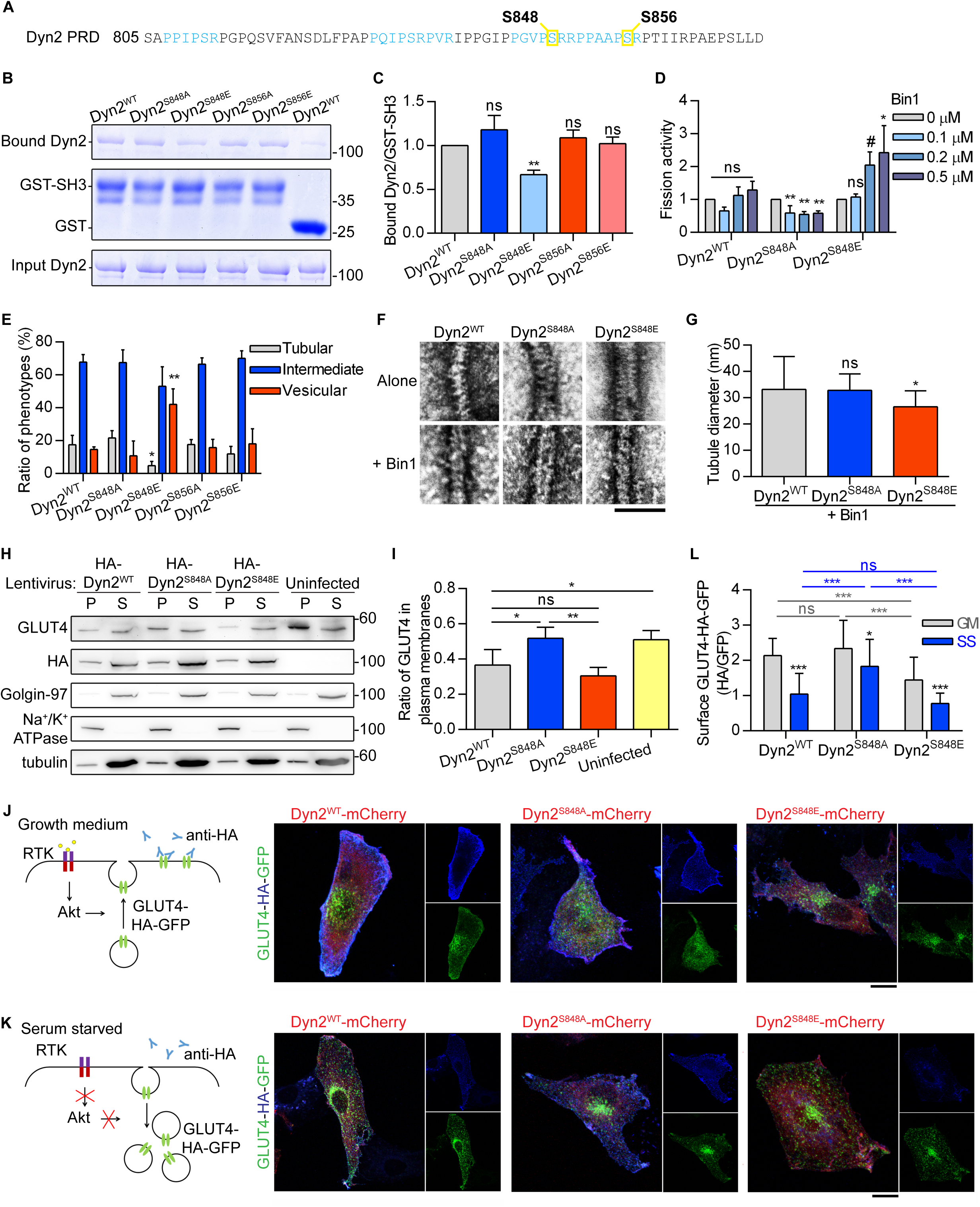
The phosphorylation of Dyn2-S848 relieves Bin1 inhibition and promotes endocytosis. (A) The amino acid sequence of Dyn2-PRD. Two potential phosphorylated serine, S848 and S856, were boxed and highlighted in yellow. The SH3-binding pockets were highlighted in light blue. (B) GST pulldown assay. GST or GST-tagged Bin1 SH3 domain was incubated with indicated Dyn2 mutants. The ratio of bound Dyn2 was detected with Coomassie Blue staining, quantified with imageJ and normalized to wild-type as shown in (C) (n = 3). (D) Fission activity of Dyn2 mutants in the presence of Bin1. Indicated Dyn2 proteins were incubated with increasing concentrations of Bin1 in the presence of GTP and SUPER templates for fission activity analysis. The data was shown as fold change relative to the Dyn2 fission activity in the absence of Bin1 (n = 3). (E) Effect of Dyn2 mutants on GFP-Bin1 tubules in myoblasts. Wild-type or indicated Dyn2-mCherry mutants were transfected into myoblasts together with Bin1-GFP. The Bin1-induced membrane morphologies were analyzed as in Fig. 3 G and shown in (E) (n ≥ 100 from three independent repeats). (F, G) Assembly of Dyn2 mutants on liposome. Indicated Dyn2 proteins were incubated with liposome alone (top) or together with Bin1 (bottom) and then visualized with negative stain TEM. Scale, 100 nm. The diameter of tubulated membrane from coincubation of Bin1 and Dyn2 were quantified and shown in (G) (n ≥ 18). (H) Subcellular distribution of endogenous GLUT4 in C2C12 myotubes expressing different Dyn2 mutants. Differentiated C2C12 myotubes were infected with lentiviruses to express indicated HA-Dyn2 then subjected to subcellular fractionation to determine the distribution of endogenous GLUT4 by Western blotting. The markers for heavy membrane (Na+/K+ ATPase, plasma membrane), light membrane (Golgin-97, trans-Golgi) and cytosol (tubulin) were used to validate this assay. The ratio of GLUT4 in heavy membrane fraction was quantified and shown in (I) (n = 4). (J-L) Effects of Dyn2-S848 mutations on the insulin-regulated GLUT4-HA-GFP distribution. L6 myoblasts co-transfected with GLUT4-HA-GFP and Dyn2-mCherry mutants were subjected to growth or serum starved media for 2 hours. The cartoons illustrate the expected location of GLUT4-HA-GFP under indicated conditions. After immunofluorescent staining with anti-HA antibody without permeabilization and imaging by confocal microscopy, the relative amount of surface GLUT4-HA-GFP was quantified by the fluorescent intensity of HA signaling in blue divided by the total GLUT4-HA-GFP in green as shown in (L). (n ≥ 27 from three independent repeats). Data are shown as average ± SD and analyzed with one-way ANOVA. ns, no significance; #P≤ 0.06; *P< 0.05; **P< 0.01; ***P< 0.001.

To measure the effect of Dyn2^S848^ phosphorylation on endocytosis in muscle cells, we monitored the internalization of transferrin and the trafficking of glucose transporter, GLUT4, for Dyn2-mediated endocytosis. With high level of endogenous Bin1 expressed in C2C12 myotubes, overexpression of Dyn2^S848A^, the phosphodeficient Dyn2 mutant, led to lower transferrin internalization and higher cell surface GLUT4 distribution than Dyn2^S848E^ and Dyn2^WT^, indicating reduced Dyn2 activity (Fig. S3 A-C and 4 H-I). Of note, the total amount of GLUT4 remained similar in cells expressing wild type or mutant Dyn2 (Fig. S3, D and E). GLUT4 membrane translocation is stimulated by insulin signaling through the phosphoinositide 3-kinase (PI3K) and Akt in muscle cells (Beg et al., 2017; Katome et al., 2003; Leto and Saltiel, 2012). To further validate the physiological importance of Dyn2-S848 phosphorylation, we utilized the L6 myoblasts to investigate the effect of Dyn2^S848^ phosphorylation on GLUT4-HA-GFP internalization triggered by serum starvation (SS). Quantification of cell surface GLUT4 by immunostaining without permeabilization showed that GLUT4 displayed a predominant intracellular distribution after three hours of SS in Dyn2^WT^-expressing cells (Fig. 4 J-L). Intriguingly, Dyn2^S848A^ overexpression resulted in significantly higher GLUT4-HA-GFP level on the cell surface upon SS, whereas Dyn2^S848E^ displayed lower surface level of GLUT4 even in the growth medium containing 10% FBS (GM), indicating that Dyn2^S848^ phosphorylation facilitates endocytosis in the muscle cells, whereas phosphodeficient Dyn2^S848A^ blocks GLUT4 internalization. Of note, neither phosphodeficient nor phosphomimetic mutants of Dyn2-Ser848 affects the exocytic pathway, as monitored by the efficiency of transferrin recycling (Fig S3, F-H).

### Phosphorylation of Dyn2-Ser848 is attenuated by insulin signaling

We next sought for evidence of phosphorylation of Dyn2-S848 by immunoprecipitation and phosphoserine detection in L6 myoblasts. The pan-phosphoserine (p-Ser) immunoblotting experiment showed significantly elevated p-Ser in precipitated HA-Dyn2^WT^ under SS, but not in HA-Dyn2^S848A^ (Fig. 5, A and B). To pinpoint the phosphorylation site of Dyn2 in a relative physiological setting, we generated a specific antibody for phosphorylated Dyn2-Ser848 (named p-S848), and detected higher p-S848 signal in C2C12 myotubes after SS (Fig. 5 C). Moreover, phosphorylation of Dyn2-S848 was significantly increased by serum starvation and reduced by insulin (Fig. 5, D and E). Similar to the effect of Dyn2^S848E^ on Bin1-GFP tubules, SS induced the fragmentation of Bin1-GFP tubules in the myoblasts, and this effect could be reversed by insulin (Fig. 5, F and G).

**Figure 5.**
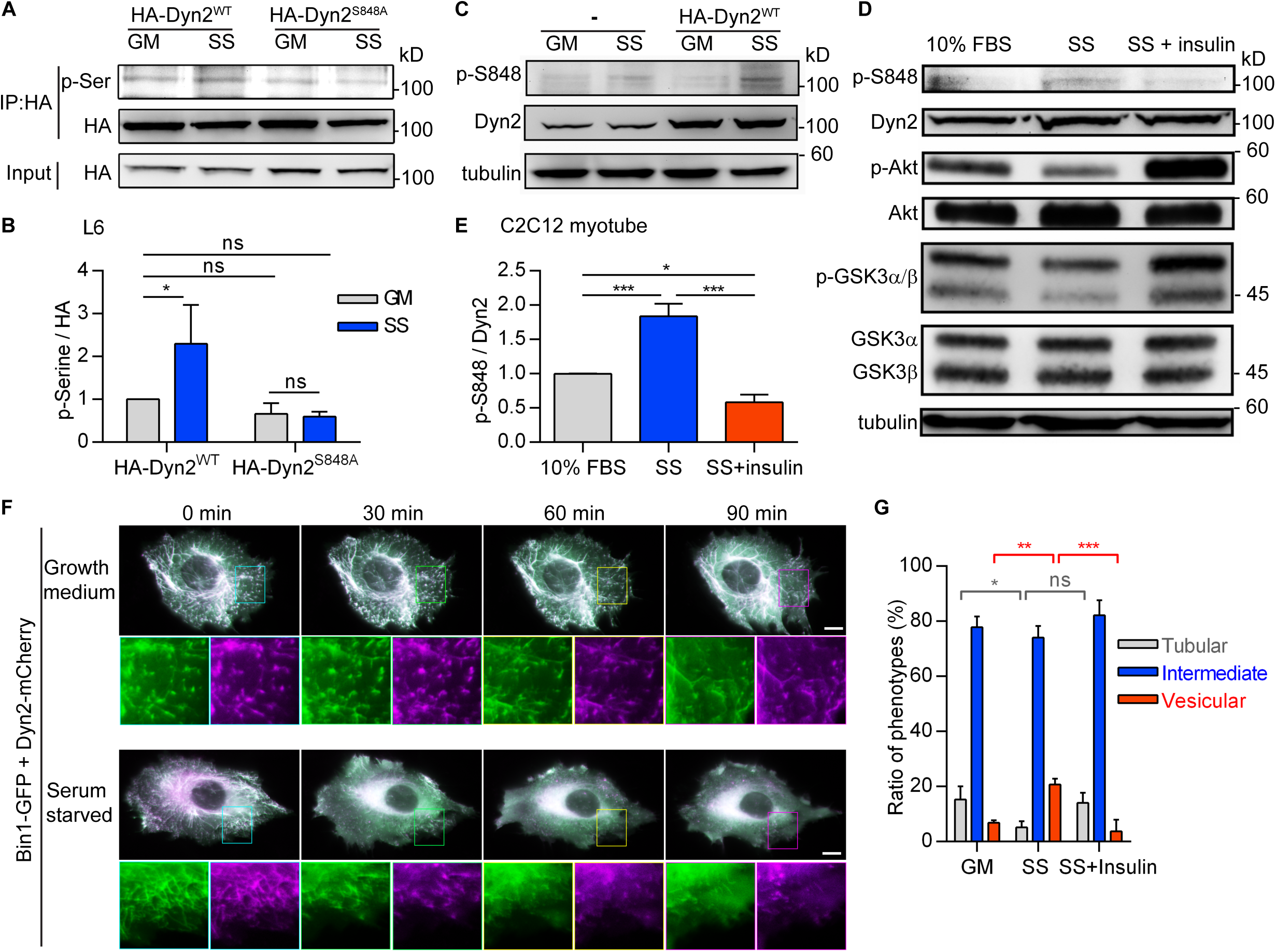
The phosphorylation of Dyn2^S848^ is regulated by insulin signaling. (A) Phosphorylation of Dyn2 in response to serum starvation. HA-tagged Dyn2^WT^ or Dyn2^S848A^ expressed in L6 myoblasts were precipitated by anti-HA antibody with or without serum starvation. Phosphorylated Dyn2 was quantified by anti-phospho-serine antibody and normalized with precipitated HA intensity, compared to HA-Dyn2^WT^ in growth medium and shown in (B) (n = 3). (C-E) Phosphorylation of endogenous Dyn2 in response to serum starvation and insulin. C2C12 myotubes were subjected to growth medium (10% FBS), 3-hour serum-free medium (SS) (C) or SS followed with 30-min, 100 nM insulin stimulation (D). Cell lysates were harvested and detected by Western blotting with indicated antibodies. The ratio of phosphorylated Dyn2^S848^ was quantified and shown in (E) (n = 3). (F) Time-lapse representative images of Bin1-GFP tubules in cells cotransfected with Dyn2-mCherry and cultured in growth or serum starved medium. Boxed areas were magnified and shown in the lower panel. Scale, 10 *μ*m. (G) Effects of insulin signaling on Bin1 tubule morphology. Bin1-GFP were transfected into myoblasts together with Dyn2-mCherry and cultured in growth, serum starved medium or starvation followed by 60 min insulin stimulation. The population of cells with tubule, intermediate or vesicular Bin1 morphology were quantified and shown in (G) (n ≥ 75 from three independent repeats). Scale, 10 *μ*m. Data are shown as average ± SD and analyzed with one-way ANOVA. ns, no significance; *P< 0.05; **P< 0.01; ***P< 0.001.

### GSK3α phosphorylates Dyn2 at Ser848

Using Scansite 4.0 to predict possible kinases of Dyn2S848, we identified glycogen synthase kinase 3α (GSK3α) as a candidate for phosphorylating Dyn2-S848 (Fig. S4 A). GSK3α is a constitutively active serine/threonine kinase that is inhibited by insulin-PI3K-Akt signaling through phosphorylation at Ser21 (Cross et al., 1995; Frame et al., 2001). We thus tested whether GSK3α increases the phosphorylation of Dyn2-Ser848 by expressing HA-Dyn2 with either wild type (WT), constitutively active (S21A, GSK3α CA), or kinase inactive (K148A, GSK3α KI) GSK3α in HeLa cells. GSK3α CA significantly enhanced p-Ser signal of HA-Dyn2^WT^ without affecting the extent of phosphorylation of HA-Dyn2^S848A^ (Fig. 6 A, B and S4 B). GSK3β, which shares high similarity in its catalytic domain with that of GSK3*α* (Woodgett, 1990), showed no effects on the phosphorylation status of S848 in HA-Dyn2^WT^and HA-Dyn2^S848A^(Fig 6, C and D), consistent with a previous report that Dyn2 is not phosphorylated by GSK3β *in vitro* (Clayton et al., 2010). We further performed an *in vitro* kinase assay and demonstrated phosphorylation of Dyn2-Ser848 by recombinant GSK3α (Fig. 6, E and F) that was confirmed by mass spectrometry detection (Fig. S4 C). Similar to the inhibitory effect of Dyn2^S848A^on transferrin endocytosis, the expression of the kinase-inactive GSK3α KI resulted in reduced transferrin internalization in the C2C12 myotubes (Fig. 6, G and H). Together, these data demonstrate that phosphorylation of Dyn2-Ser848 is a critical regulatory target for Bin1-SH3 and GSK3*α* that tunes the membrane fission functions of Dyn2 to control endocytosis in muscle cells.

**Figure 6.**
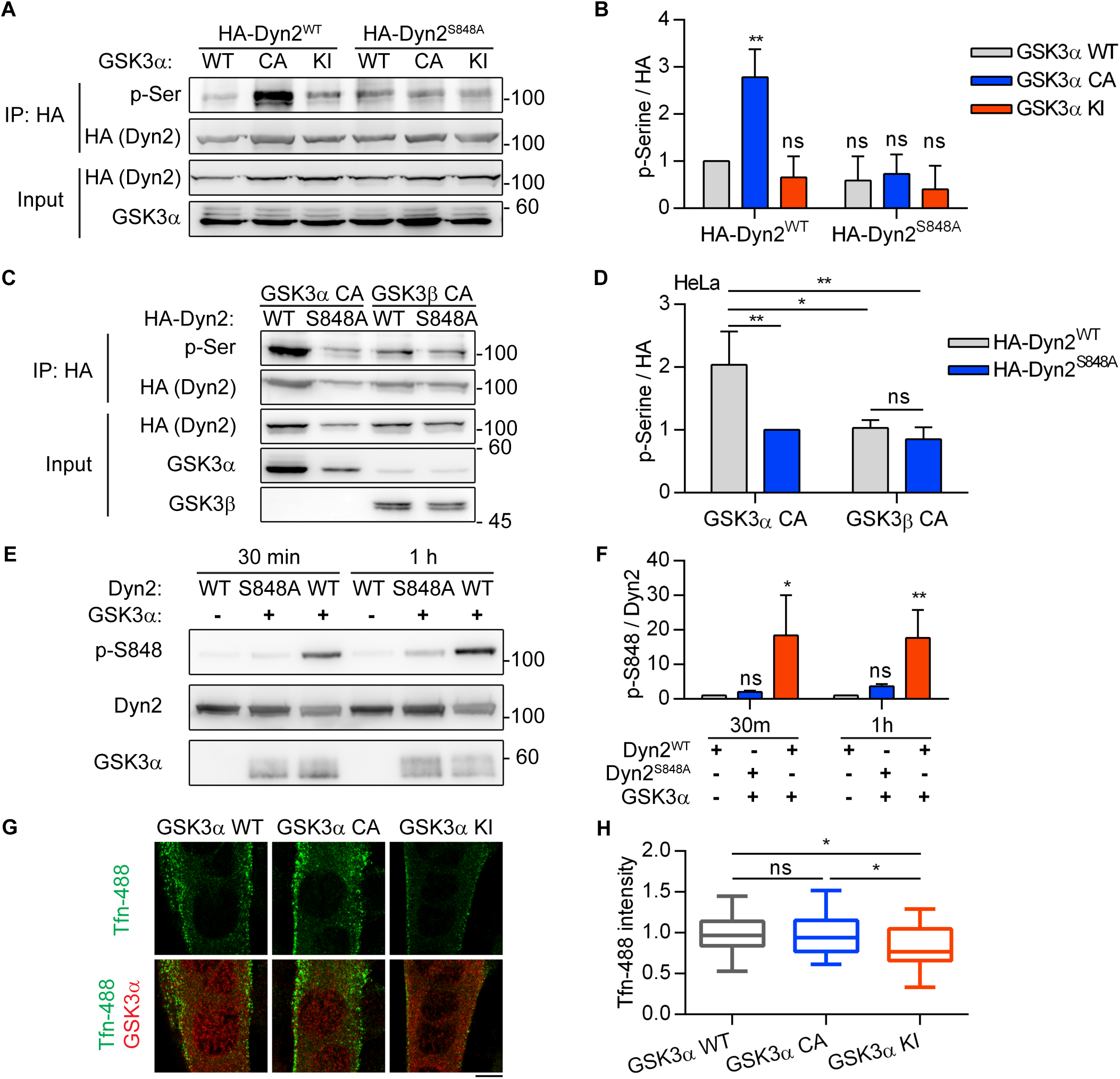
GSK3*α* phosphorylates the S848 residue of Dyn2. (A) Phosphorylation of Dyn2 in response to GSK3*α* overexpression. HA-Dyn2^WT^ or HA-Dyn2^S848A^ were co-expressed with WT, CA or KI forms of GSK3*α* in Hela cells. After precipitation with anti-HA antibody, the phosphorylated Dyn2 was detected with anti-phospho-serine antibody, normalized with HA intensity and shown in (B) (n = 3). (C) Isoform specific activity of GSK3 on Dyn2. HA-Dyn2^WT^ or HA-Dyn2^S848A^ were co-expressed with constitutive active (CA) GSK3*α* or GSK3α in Hela cells. After precipitation with anti-HA antibody, the phosphorylated Dyn2 was detected with anti-phospho-serine antibody, normalized with HA intensity and shown in (D) (n = 3). (E) *In vitro* kinase assay. 0.8 *μ*g purified Dyn2^WT^or Dyn2^S848A^were incubated with ATP in the presence or absence of 10 ng purified GSK3*α*. After incubation 30 min or 1 hour incubation, the phosphorylated Dyn2 was quantified by Western blotting with anti phospho-Dyn2^S848^antibody. The intensity of p-Dyn2^S848^was quantified with ImageJ, normalized with Dyn2 signal and shown in (F) (n = 3). (G) Effect of GSK3*α* mutants on transferrin internalization in myotubes. C2C12 myoblasts transfected with indicated GSK3*α* mutants and differentiated into myotubes were incubated with Alexa-488 labelled transferrin at 37°C for 20 min. Cells were then washed on ice, fixed and imaged under confocal microscopy. Scale, 10 µm. Fluorescence intensity of internalized transferrin was quantified and shown in (H) (n = 25 from three independent repeats). Bar graphs are shown as average ± SD, box plots are shown as median ± min and max value. Data are analyzed with one-way ANOVA. ns, no significance; *P< 0.05; **P< 0.01. (I)

## DISCUSSION

In this work, we uncover Dyn2-S848 as a target for regulating Dyn2 activity by GSK3*α* and Bin1-SH3, and the phosphorylation status of Dyn2-S848 is a central switch that shapes Dyn2-Bin1 interaction from antagonism to synergy for membrane fission and endocytosis (Fig. 7). Increased activity of Dyn2 in CNM-Bin1-expressing myoblasts with fragmented tubules is reminiscent of disrupted T-tubule network in mice expressing mutant Bin1 that lacks the SH3 domain (Fig. 3 F) (Silva-Rojas et al., 2021). On the other hand, CNM-associated Dyn2 mutants display enhanced fission activity as a result of lacking auto-inhibition, which leads to disrupted T-tubules in *Drosophila* and mouse muscles (Chin et al., 2015; Cowling et al., 2011). The phenotypic similarities between muscles that express CNM-Bin1 and CNM-Dyn2, and the tunable Dyn2-Bin1 interaction identified in this study suggest that Dyn2 hyperactivity is a common pathogenic mechanism of CNM. In line with that, both Dyn2 depletion and Bin1 overexpression could rescue CNM phenotype in several CNM mouse models, including myotubularin (MTM1) (Cowling et al., 2014; Lionello et al., 2019; Tasfaout et al., 2017). MTM1 is a phosphoinositide 3-phosphatase, whose mutations lead to the X-linked form of CNM (Laporte et al., 2000; Laporte et al., 1996). MTM1 has been previously identified as a binding partner of Bin1 and enhancing Bin1-mediated membrane tubulation (Royer et al., 2013). It is possible that MTM1 could also function as a negative regulator of Dyn2 either directly or indirectly through Bin1. Further study on how these three proteins physically and functionally interact in skeletal muscle would bring more insights on the pathological progression of CNM.

**Figure 7.**
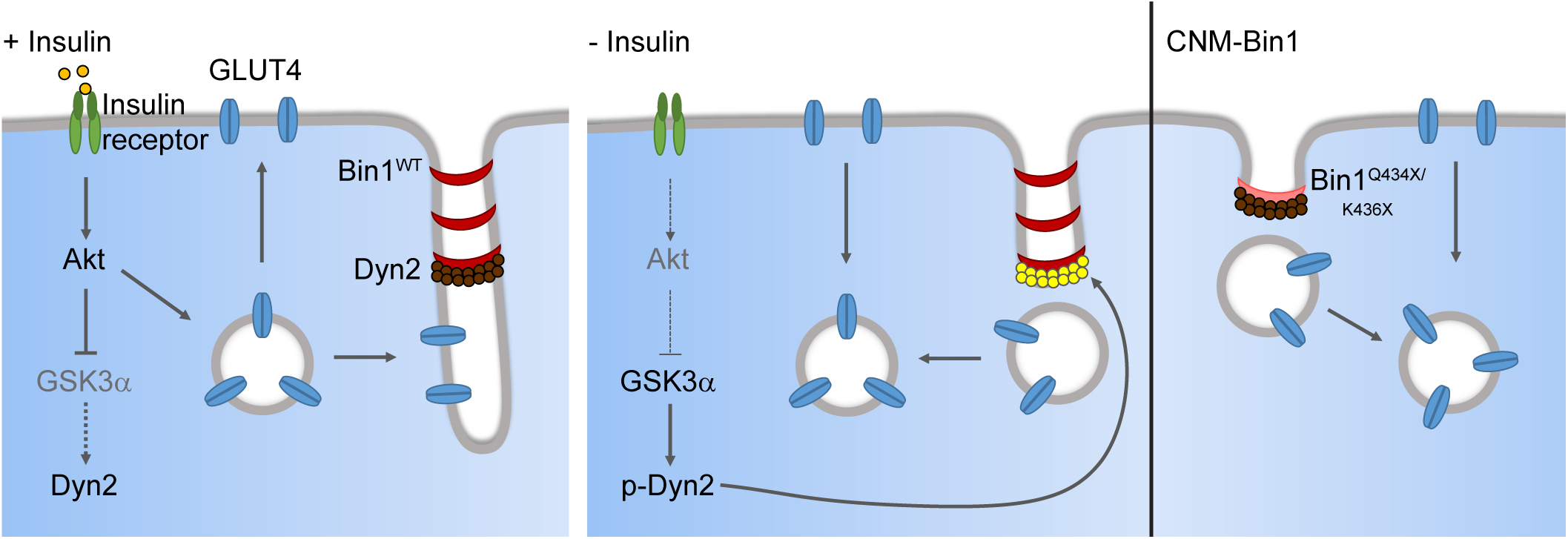
Dyn2-S848 phosphorylation serves as a molecular switch to promote endocytosis in muscle cells. Dyn2 activity is inhibited by the binding of Bin1 while GSK3α is inactivated by external signaling, e.g. insulin-PI3K-Akt. The attenuated endocytosis together with the increased exocytosis of GLUT4 lead to its efficient plasma membrane translocation and glucose uptake of muscle upon insulin stimulation. After insulin signaling turns off, GSK3α becomes active and phosphorylates Dyn2-S848 to relieve the Bin1 inhibition and promote endocytosis. The internalization of GLUT4 from muscle surface ceases the glucose uptake of skeletal muscle. CNM-related Bin1 mutations with partial truncation of the SH3 domain lose their inhibitory effect on Dyn2 thus result in hyperactive Dyn2 and fragmented T-tubule in muscle cells.

The location and function of dynamin is regulated by its binding partners, which may positively or negatively regulate its activity (Hohendahl et al., 2017; Meinecke et al., 2013; Neumann and Schmid, 2013). Here, we find that Dyn2-S848 phosphorylation reduces Bin1 interaction without affecting its binding to endophilin-SH3, indicating endophilin A1 and Bin1 do not bind to the same region of Dyn2, which is in agreement with previous study (Solomaha et al., 2005). As a result, the SH3 domains from Bin1 and endophilin have vastly different effects on the activity of Dyn2 (Fig. 1 K). Given the differential tissue expression of endophilin A1 and Bin1 which is neuron-specific and ubiquitously expressed, respectively (Lee et al., 2002; Ringstad et al., 1997), we speculate that the activity of Dyn2 is highly demanded in neuron whereas it needs to be attenuated and tightly controlled in muscle. The multi-layered regulation of Dyn2 activity for endocytosis identified here warrants detailed structure-function analysis of the interactions between Dyn2 and its binding proteins for the regulation of endocytosis and membrane remodeling in different tissues.

The conventional dynamin is believed to evolve as a central regulator for the late stage of endocytosis in holozoans, which includes animals and their closest single-celled relatives but not fungi (Dergai et al., 2016; Liu et al., 2012). Although several regulatory mechanisms have been identified to control the activity of neuronal dynamin-1 (Armbruster et al., 2013; Clayton et al., 2010; Smillie and Cousin, 2005), little is known regarding the regulation of Dyn2 activity upon cellular demands (Ahn et al., 2002). Here the identification of GSK3*α* as a critical kinase that relieves Dyn2 from the inhibition of Bin1-SH3 provides a conceptual framework that could foster future discovery of other posttranslational modifications that fine-tune the activity of endocytic proteins to meet the need of various cellular milieu. Given the extracellular signal-regulated kinase 1 (Erk1), a component of the mitogen-activated protein kinase (MAPK) pathway, is also predicted as a potential kinase that phosphorylates Dyn2-S848 (Fig. S4 A), it will be of great interest to investigate the potential function of Erk1 on Dyn2 activity and the effect of mitogen signaling on endocytosis regulation.

## MATERIALS AND METHODS

### Protein expression and purification

Human dynamin-2 isoform 1 were expressed in Sf9 insect cells transiently transfected with various constructs and purified via affinity chromatography using the SH3 domain of amphiphysin-2 as reported previously (Liu et al., 2011b). Various constructs of human Bin1 isoform 8 were cloned into pGEX-4T-1 or pET30a vectors for expression in *E. coli* and purified with glutathione Sepharose beads (GE) or Ni-NTA spin columns (Qiagen), respectively, followed by elution as recommended by the manufacturer. Mouse endophilin A1 was purified like Bin1.

### Preparation of lipid templates

Membrane compositions used in this study were DOPC:DOPS:PI(4,5)P_2_ at 80:15:5 or DOPC:DOPS:PI(4,5)P2:Rh-PE at 79:15:5:1 for fluorescent-tagged membrane. Liposomes and SUPER templates were prepared as previously described (Neumann et al., 2013). Briefly, for liposome preparation, lipid mixtures were dried, rehydrated in buffer containing 20 mM HEPES (pH 7.5) and 150 mM KCl then subjected to three rapid freeze-thaw cycles followed by extrusion through a 0.1 µm polycarbonate membrane (Whatman) using Avanti mini extruder. SUPER templates were generated by incubating 2.5 µm silica beads in a solution containing 100 µM fluorescent-tagged liposomes and 1 M NaCl for 30 min at room temperature. Excess unbound liposomes were washed four times with water after incubation.

### Fluorescence microscopy

For SUPER templates visualization, images were collected using an inverted Zeiss Axio Observer Z1 microscope equipped with plan-apochromat 100 ×/1.40 oil DIC M27 objective (Carl Zeiss) and analyzed using ZEN software (Carl Zeiss). For cells fixed with 4% formaldehyde and permeabilized with 0.1% saponin, images were acquired using a confocal microscope (LSM700, Carl Zeiss) equipped with plan-apochromat 63×/1.40 oil DIC M27 objective (Carl Zeiss) and processed using ZEN software (Carl Zeiss).

### Fission and SUPER template tubulation assays

Fission activity of Dyn2 was measured as previously described (Chin et al., 2015). Briefly, SUPER templates were incubated with 0.5 µM Dyn2, 1 mM GTP and indicated concentrations of Bin1 variants in assay buffer (20 mM HEPES (pH 7.5), 150 mM KCl, and 1 mM MgCl_2_) for 30 min at room temperature. The mixtures were spun down at low speed (260 x *g*) in swinging bucket rotor to separate SUPER template from the released vesicles. Fluorescence intensity was measured in a plate reader (RhPE excitation = 530/25 nm bandwidth and emission = 580/25 nm bandwidth). The fission activity was expressed as the fraction of released lipids from total SUPER templates.

For visualization of membrane tubulation, SUPER template were slowly added in an evenly distributed manner to solution containing 20 mM HEPES (pH 7.5), 150 mM KCl and 0.5 µM wild-type or mutant Bin1 on BSA coated eight-chamber slides. After 10 min incubation at room temperature, images of SUPER templates were acquired using Zeiss inverted fluorescent microscope Axio Observer Z1 with a 100×/1.40-NA oil-immersion objective with CCD monochrome Photometrics CoolSNAP HQ.

### Liposome binding, transmission electron microscopy (TEM) and GTPase activity assays

To analyze membrane binding ability, 0.5 µM Dyn2 and/or Bin1 were incubated with 150 µM, 100 nm liposomes at 37°C for 10 min. Pellets containing liposome-bound proteins were separated from soluble fractions containing unbound proteins (supernatant) by centrifugation at 15,000 x *g* and resuspended in assay buffer (20 mM HEPES (pH 7.5), 150 mM KCl). Pellet and supernatant were run on SDS-PAGE, stained with Coomassie Blue and protein intensities were quantified using ImageJ. Membrane binding ability was expressed as the fraction of membrane-bound proteins divided by total proteins.

To visualize protein assembly on membrane, 1 µM Dyn2 and/or 1 µM Bin1 were incubated with 25 µM liposome for 10 min at 37°C. The mixture was then adsorbed onto the surface of glow-discharged (30 s), carbon film-supported grids (200 mesh copper) for 5 min followed by 2 min negative-staining with 2% uranyl acetate. Images were captured with a Hitachi H-7650 electron microscope operated at 75 kV and a nominal magnification of 150,000x. Liposome tubules diameter were measured using ImageJ.

For GTPase activity assay, 0.5 µM purified Dyn2 was incubated with 0.5 µM Bin1 or Endophilin in reaction mixture containing 150 µM liposomes, 20 mM HEPES (pH 7.4), 150 mM KCl, 1 µM MgCl_2_ and 1mM GTP at 37°C. Liposome-stimulated GTPase activity of Dyn2 was measured as a function of time using a colorimetric malachite green assay which monitors the release of inorganic phosphate (Leonard et al., 2005).

### GST pulldown assay

To analyze SH3-PRD binding affinity, purified His-Bin1 proteins were first incubated with 100 nm liposomes containing 5% PI(4,5)P_2_ to alleviate the autoinhibition between SH3 domain and PI motif before being incubated with GST or GST-tagged PRD of Dyn2 immobilized on glutathione beads for 30 min at room temperature. After washing, the beads were boiled in sample buffer and bound protein were detected by SDS-PAGE followed by Coomassie Blue staining. ImageJ software was used to quantify protein bands intensity of bound Bin1 relative to GST-PRD. Pull down assay for purified Dyn2 mutants with GST-tagged SH3 domain of Bin1 or endophilin was performed in a similar manner without the presence of 100 nm liposomes.

### Cell culture, transfection and lentiviral infection

Mouse-derived C2C12 myoblasts (ATCC, CRL-1772) were cultured in growth medium (GM) composed of high glucose (4.5 mg/mL) DMEM supplemented with 2mM _L_-glutamine, 1 mM sodium pyruvate, antibiotics, and 10% fetal bovine serum (Gibco). Upon reaching 90% confluency, C2C12 differentiation was induced by replacing GM with differentiation medium (DM) composed of high glucose (4.5 mg/mL) DMEM supplemented with 1 mM sodium pyruvate, antibiotics, and 2% horse serum (Gibco). The first time cells were incubated in DM was regarded as day 0 of differentiation. Rat-derived L6 myoblasts (ATCC, CRL-1458) were cultured in low glucose (1.0 mg/mL) DMEM supplemented with antibiotics and 10% fetal bovine serum (Gibco). For transfection, cells at 70% confluency were transfected with plasmid of interest using Lipofectamine 2000 (Invitrogen) or *Trans*IT-X2 (Mirus Bio) as suggested by the manufacturers. For lentiviral infection, C2C12 myoblasts at 50% confluency were infected with viruses in the presence of 12 µg/ml polybrene followed by puromycin (2µg/ml) selection 24h post-infection for 3 days before DM (containing 2% FBS) replacement. 10 µg/ml doxycycline were added to day 3 differentiated C2C12 and GLUT4 distribution in cells was analyzed at day 6 of differentiation. List of plasmids used for transfection and lentiviral generation in this study could be found in Table S1.

### Live cell imaging

For time-lapse microscopy, cells were imaged with spinning disc confocal microscope or lattice-light sheet microscope (Chen et al., 2014). For spinning disc confocal microscope, C2C12 myoblasts were seeded on glass-bottom dishes and cotransfected with Bin1-GFP and Dyn2-mCherry. Cells were placed at 37°C in imaging medium (phenol-red free DM with 20 mM HEPES, pH 7.4, 50 μg/ml ascorbic acid, and 10% FBS) and images were captured with 200 ms (488 nm laser) and 400 ms (561 nm laser) exposure and 5 s interval time using Carl Zeiss Cell Observer SD equipped with plan-apochromat 100×/1.40 oil DIC M27 objective (Carl Zeiss). For lattice light-sheet microscopy, C2C12 myoblasts cotransfected with Bin1-GFP and Dyn2-mCherry were immersed in an imaging medium filled chamber. Images were illuminated by exciting each plane with a 488 nm laser at 12.56 nano-Watt (nW) (at the back aperture of the excitation objective) and 561 nm laser at 37.2 nW, with an excitation outer/inner numerical aperture parameters of 0.55/0.44. Orthogonal to the illumination plane, the water immersed objective lens (Nikon, CFI Apo LWD 25XW, 1.1 NA, 2 mm WD) mounted on a piezo scanner (Physik Instrumente, P-726 PIFOC) is used to collect the fluorescence signal, which is then imaged through an emission filter (Semrock Filter: FF01-523/610-25) onto sCMOS camera (Hamamatsu, Orca Flash 4.0 v2 sCOMS) through a 500 mm tube lens (Edmund 49-290, 500 mm FL/50 mm dia; Tube Lens/TL) to provide the 63X magnification observation. And then the cells were imaged through entire cell volumes at 5 s intervals by using the sample piezo (Physik Instrumente, P-622 1CD) scanning mode. Finally, raw data were then deskewed and deconvoluted using GPU_decon_bin (Chen et al., 2014). And using Amira software (Thermo Fisher) to display the 3D images.

### Transferrin uptake assay

Transferrin uptake assay was performed as previously reported (Liu et al., 2008). Briefly, C2C12 transfected with mCherry-tagged Dyn2 mutants and seeded on coverslips were incubated with 5 μg/ml Alexa-488-conjugated transferrin for 10 min (myoblasts) or 20 min (myotubes) at 37°C. For recycling efficiency, 30 min incubation with transferrin-488 was followed by 30 min incubation in growth medium without transferrin-488. Cells were washed with ice-cold PBS then with acid buffer (150 mMNaCl and 150 mM glycine, pH 2.0) repeatedly. Afterward cells were fixed with 4% FA and mounted on glass slides using Fluoromount-G mounting medium (Southern Biotech). Images of the cells were acquired using confocal microscopy and analyzed with ZEN software (Carl Zeiss).

### Subcellular fractionation and analysis of surface:total GLUT4 ratio

Subcellular fractionation to analyze GLUT4 distribution was done as previously described (Laiman and Liu, 2020). Briefly, lentiviral infected C2C12 myotubes were homogenized in ice-cold HES-PI buffer (255 mM sucrose, 20 mM HEPES (pH 7.4), 1 mM EDTA, cOmplete protease inhibitors (Roche)). The lysates were then cleared by centrifugation at 1,000 x *g* for 5 min at 4°C. After centrifugation at 16,000 x *g* for 20 min at 4°C, the supernatant was separated from pellet and subjected to TCA precipitation for 1 h at 4°C. Both pellet and supernatant were loaded to SDS-PAGE followed by immunoblotting to detect GLUT4 and markers for heavy membrane, light membrane and cytosol. All antibodies used in this study are listed in Table S2.

Insulin-stimulated HA-GLUT4-GFP localization was done according to previous report (Zeigerer, Lampson et al., 2002). Briefly, L6 were seeded on fibronectin coated coverslips then cotransfected with Dyn2 mCherry and GLUT4-HA-GFP. Cells were incubated in growth medium or serum starved for 2 h prior to fixation and subjected to immunofluorescent staining with anti HA without permeabilization. Cell was imaged under LSM700 confocal microscope (Carl Zeiss) equipped with plan-apochromat 63×/1.40 oil DIC M27 objective (Carl Zeiss) and analyzed using ZEN software (Carl Zeiss). The relative amount of surface GLUT4-HA-GFP was quantified by the fluorescent intensity of HA signal divided by the total GFP signal.

### Immunoprecipitation

For immunoprecipitation, cells were washed with ice-cold 1x PBS added with 1.5 mM Na3VO4 and 50mM NaF before being lysed in lysis buffer (1X PBS, 1% Tx-100, 1.5 mM Na3VO4, 50mM NaF, PhosSTOP (Roche) and cOmplete protease inhibitors (Roche)). The lysates were centrifuged at 16,000 x *g* for 10 min at 4°C. The supernatant was then incubated with anti-HA Agarose (Sigma-Aldrich, Cat# A2095) for 1 h at 4°C on a rotator. The beads were washed before being boiled in sample buffer for the following SDS-PAGE and immunoblotting. All antibodies used in this study are listed in Table S2. Band intensities were quantified using ImageJ.

### *In vitro* kinase assay

For in vitro phosphorylation of Dyn2, recombinant Dyn2 was purified from Sf9 as mentioned above, active recombinant GSK3α was purchased (Millipore, Cat# 14-492). 0.8 µg Dyn2WT or Dyn2S848A was incubated with 10 ng GSK3α in kinase buffer (50 mM HEPES (pH 7.4), 15 mM MgCl2, 200 µM Na3VO4, 100 µM ATP). The total reaction volume was 30 µl. Following 30 min or 1 h incubation at 30°C, reaction was stopped by adding sample buffer and was boiled at 90°C before subjected to SDS-PAGE and Western blot analysis with phospho-S848-specific (p-S848) antibody. p-S848 antibody is homemade polyclonal antibody through immunization against phosphoSer848 on Dyn2 using synthetic phosphopeptide with following sequence: 844 PGVP(pSer)RRPPAAPSRC 858, and purified with affinity column.

### Mass spectrometry

For analysis of Dyn2 phosphorylation status, recombinant Dyn2 phosphorylated by GSK3α in vitro as mentioned above was subjected to SDS-PAGE. The gel was then stained with Coomassie Blue and bands containing Dyn2 were cut from the gels as closely as possible. The gel pieces were destained, followed by reduction, alkylation and dehydration before undergoing in-gel digestion with trypsin. The resulting peptides were extracted and went through desalting in C18 column before LC-MS/MS analysis. Peptides were separated on an UltiMate 3000 LCnano system (Thermo Fisher Scientific). Peptide mixtures were loaded onto a 75 μm ID, 25 cm length C18 Acclaim PepMap NanoLC column (Thermo Scientific) packed with 2 μm particles with a pore of 100 Å. Mobile phase A was 0.1% formic acid in water, and mobile phase B was composed of 100% acetonitrile with 0.1% formic acid. A segmented gradient in 50 min from 2% to 40% solvent B at a flow rate of 300 nl/min. LC-MS/MS analysis was performed on an Orbitrap Fusion Lumos Tribrid quadrupole mass spectrometer (Thermo Fisher Scientific). Targeted mass spectrometry analysis was performed. Mass accuracy of <5 ppm, and a resolution of 120,000 at m/z=200, AGC target 5e5, maximum injection time of 50 msec) followed by HCD-MS/MS of the focused on m/z 633.8338 (2+) and m/z 422.8916 (3+). High-energy collision activated dissociation (HCD)-MS/MS (resolution of 15,000) was used to fragment multiply charged ions within a 1.4 Da isolation window at a normalized collision energy of 32. AGC target 5e4 was set for MS/MS analysis with previously selected ions dynamically excluded for 180 s. The MS/MS spectra of pS848 phosphopeptide were manually identified and checked.

### Statistical analysis

Quantitative data in this study are expressed as mean ± SD from at least three independent experiments. GraphPad Prism 8.0 was used for statistical analysis and graphs generation. All data were analyzed with one-way ANOVA followed by Dunnett’s or Tukey’s *post hoc* test, or Student’s t test. Statistical significance was defined using GraphPad Prism 8.0. P < 0.05 was considered statistically significant, indicated as *P< 0.05; **P< 0.01; ***P< 0.001.

### Data Availability

Uncropped gels, blots and all data sets are included as source data.

## Acknowledgement

We are extremely grateful to Dr. Chun-Liang Pan (NTU) and Dr. Sandra Schmid (Chan Zuckerberg Biohub) for critical reading and helpful comments on this paper. We thank Dr. Timothy E. McGraw for sharing HA-GLUT4-GFP plasmid, and the mass spectrometry technical research services from NTU Consortia of Key Technologies and NTU Instrumentation Center, the imaging core at the First Core Labs, as well as the electron microscopy (EM) core in NTU for their technical support. This work was supported by Ministry of Science and Technology grants MOST 109-2628-B-002-051 and National Taiwan University Hospital (NTUH) translational medicine grant to Y.-W. Liu.

## Author contribution

All authors participated in the experimental design; J. Laiman, J. Loh, MC and YL performed major experiments and analyzed data; WT and BC assisted in live cell imaging and data interpretation; J. Laiman and YL wrote the manuscript; BC, YC, LC and YL supervised the project.

The authors declare no competing interest.

## Supplementary Figures

**Fig S1.**
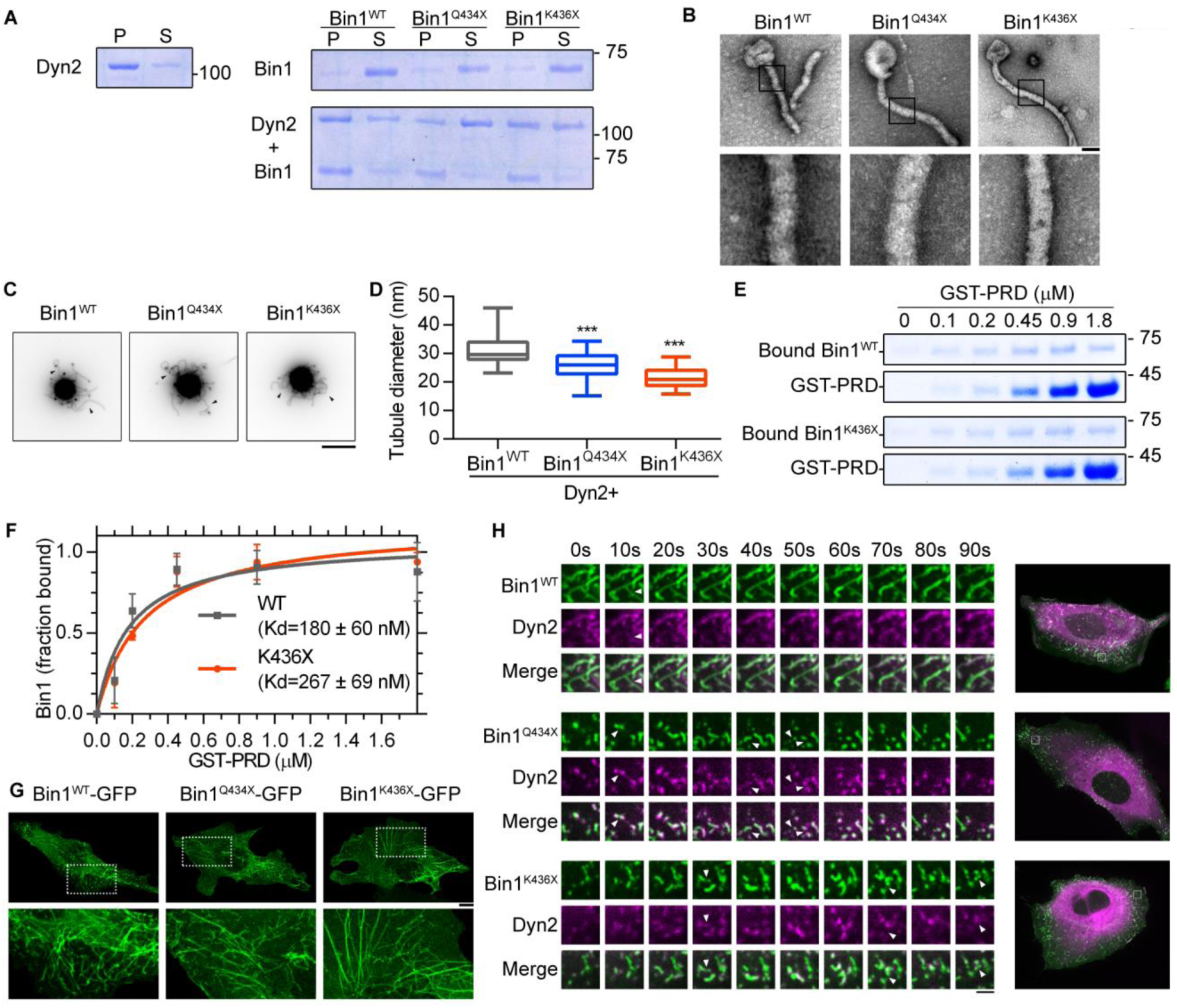
(related to Fig 3) Biochemical properties of CNM-related Bin1 mutants with SH3 domain truncation. (A) Membrane binding ability of CNM-Bin1-SH3 mutants. 0.5 µM Bin1 mutants and/or 0.5 µM Dyn2 were incubated with 100 nm liposome for 10 min at 37°C. Liposome-bound proteins (pellet, p) were separated from unbound ones (supernatant, s) through centrifugation sedimentation. (B) Assembly of Bin1 mutants on liposome. 1 µM Bin1 were incubated with liposome and then visualized with negative stain TEM. Scale, 100 nm. Boxed areas were magnified and shown as below. (C) Membrane tubulation ability of Bin1 mutants. 0.5 µM wild-type and mutant Bin1 were incubated with SUPER templates for 10 min at room temperature and imaged under fluorescent microscopy. Black arrow heads indicate the tubulated membrane. Scale, 5 µm. (D) Quantification of tubulated liposome diameter from Bin1 and Dyn2 coincubation for Fig. 3 E shown as median ± min and max value (n = 21). (E-F) Binding affinity between Bin1 and the PRD of Dyn2. Increasing concentrations of GST-PRD was used to pull down 0.2 µM purified Bin1^WT^ or Bin1^K436X^, PRD- bound Bin1 was then analyzed. The dissociation constant (Kd) was determined by curve fitting using nonlinear regression and shown with SD in (F) (n = 3). (G) Membrane tubulation ability of Bin1 mutants in cellulo. GFP-tagged Bin1 mutants were over-expressed in C2C12 myoblasts through transfection and imaged under confocal microscopy. Scale, 10 µm. (H) Time-lapse representative images of CNM-associated Bin1 mutants co-expressed with Dyn2-mCherry in C2C12 myoblast were magnified and shown. White arrow heads indicate the occurrence of membrane fission. Scale, 2 µm. Data are analyzed with one-way ANOVA or Student’s t-test. ns, no significance; *P< 0.05; ***P< 0.001.

**Fig S2.**
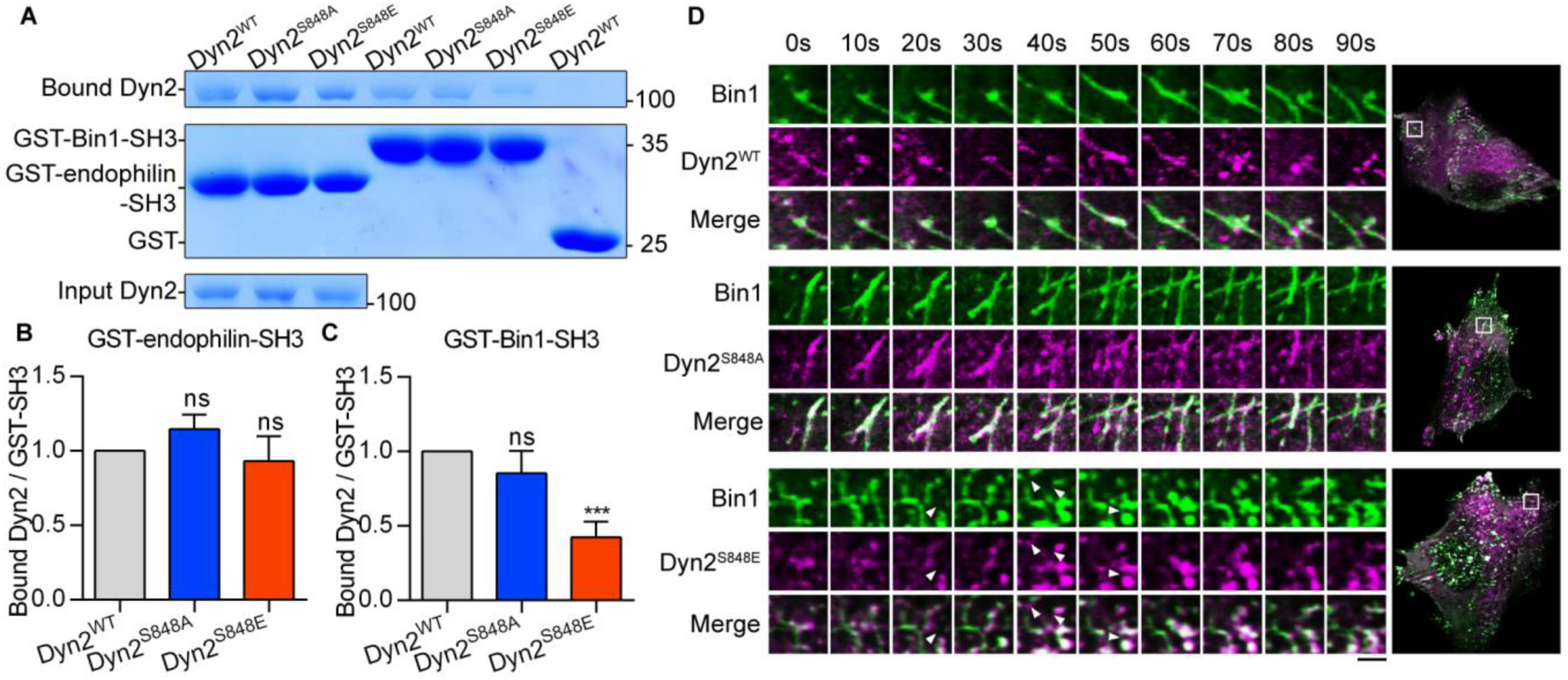
(related to Fig 4) Dyn2-Ser848 phosphorylation relieves Bin1 inhibition and promotes membrane fission. (A-C) Binding ability between Dyn2 and different SH3 domain. 12 µg GST, GST-Bin-SH3 or GST-Endophilin-SH3 was incubated with 6µg indicated purified Dyn2. Bound Dyn2 was analyzed with SDS-PAGE and Coomassie Blue staining. The ratio of bound Dyn2 was quantified, normalized to wild-type and shown in (B) and (C) (n = 3). (D) Time-lapse representative images of Bin1-GFP tubules in cells co-expressing wild-type, S848A or S848E Dyn2-mCherry were magnified and shown. White arrow heads indicate the occurrence of membrane fission. Scale, 2 µm. Bar graphs are shown as average ± SD, box plots are shown as median ± min and max value. Data are analyzed with one-way ANOVA. ns, no significance; *P< 0.05; ***P< 0.001.

**Fig S3.**
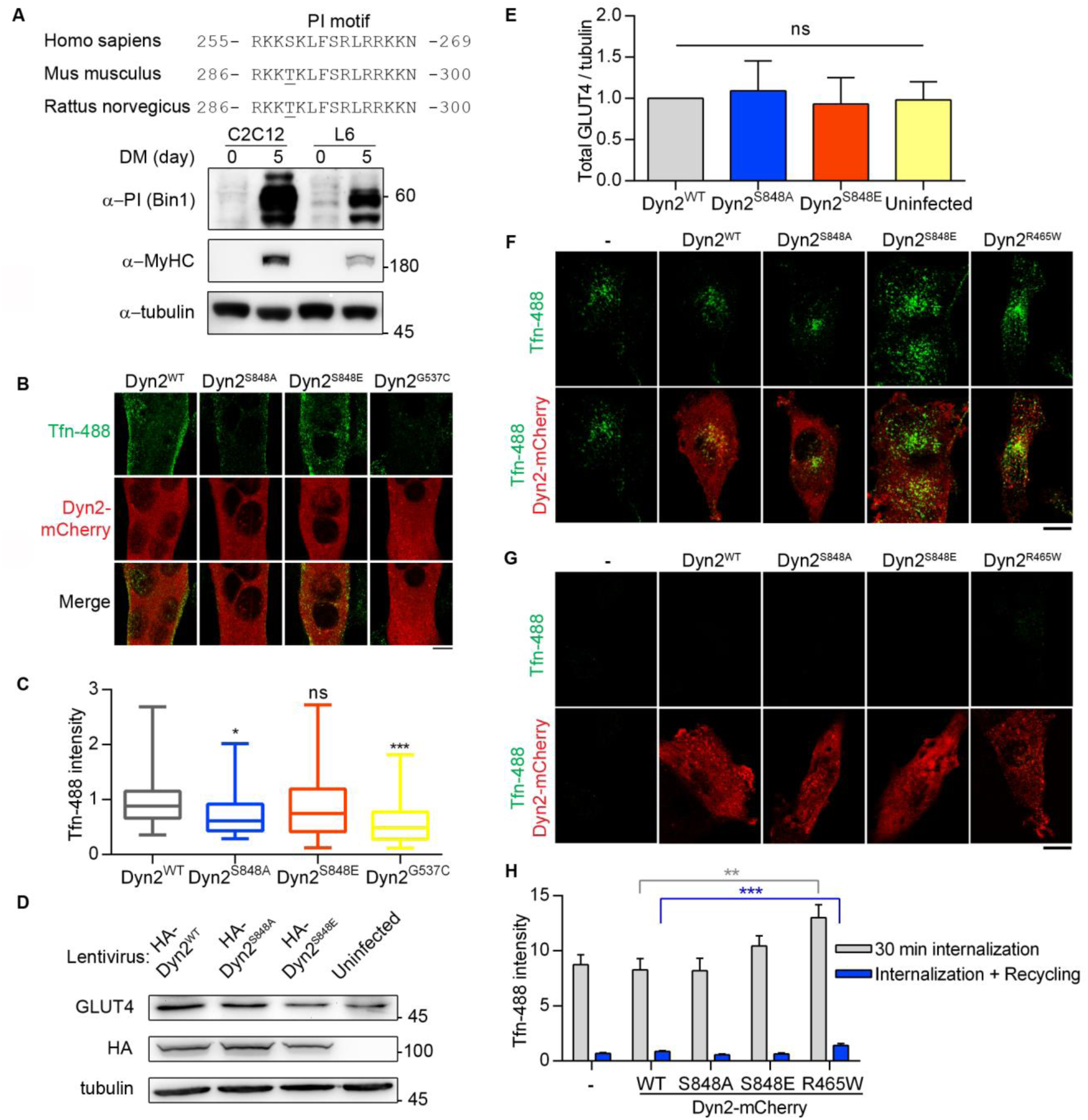
(related to Fig 4) Dyn2-Ser848 phosphorylation promotes endocytosis, but not exocytosis. (A) Amino acid sequences of Bin1-PI motif from different organisms (top). The expression of muscle specific Bin1 in C2C12 or L6 myoblasts cultured in growth or differentiation medium was examined by Western blotting using anti-PI motif antibody (bottom). (B-C) Effect of Dyn2 mutants on transferrin internalization in myotubes. C2C12 myoblasts transfected with indicated Dyn2-mCherry mutants and differentiated into myotubes were incubated with Alexa-488 labelled transferrin at 37°C for 20 min. Cells were then washed on ice, fixed and imaged under confocal microscopy. Scale, 10 µm. Fluorescence intensity of internalized transferrin was quantified, normalized to Tfn-488 intensity of Dyn2^WT^transfected cells and shown in (C) (n ≥ 24 over two independent repeats). (D-E) The amount of endogenous GLUT4 in C2C12 myotubes expressing different Dyn2 mutants. C2C12 myotubes infected with indicative HA-Dyn2 mutations lentiviruses expressed similar level of GLUT4, and the data was quantified, normalized to wild-type and shown in (E) (n = 4). (F-H) Effects of Dyn2-S848 mutants on transferrin internalization or recycling in L6 myoblast. Transferrin uptake of L6 myoblasts, L6 expressing indicative mutant Dyn2-mcherry was subjected to 30-min transferrin-488 uptake, wash and imaging (F). The recycling efficiency of these internalized transferrin was analyzed by chasing the signal by 30-min incubation of growth medium without trasnferrin-488 (G). The intensity of internalized transferrin-488 was quantified and shown in (H) (n ≥ 25). Scale, 10 µm. Bar graphs are shown as average ± SD, box plots are shown as median ± min and max value. Data are analyzed with one-way ANOVA. ns, no significance; *P< 0.05; ***P< 0.001.

**Fig S4.**
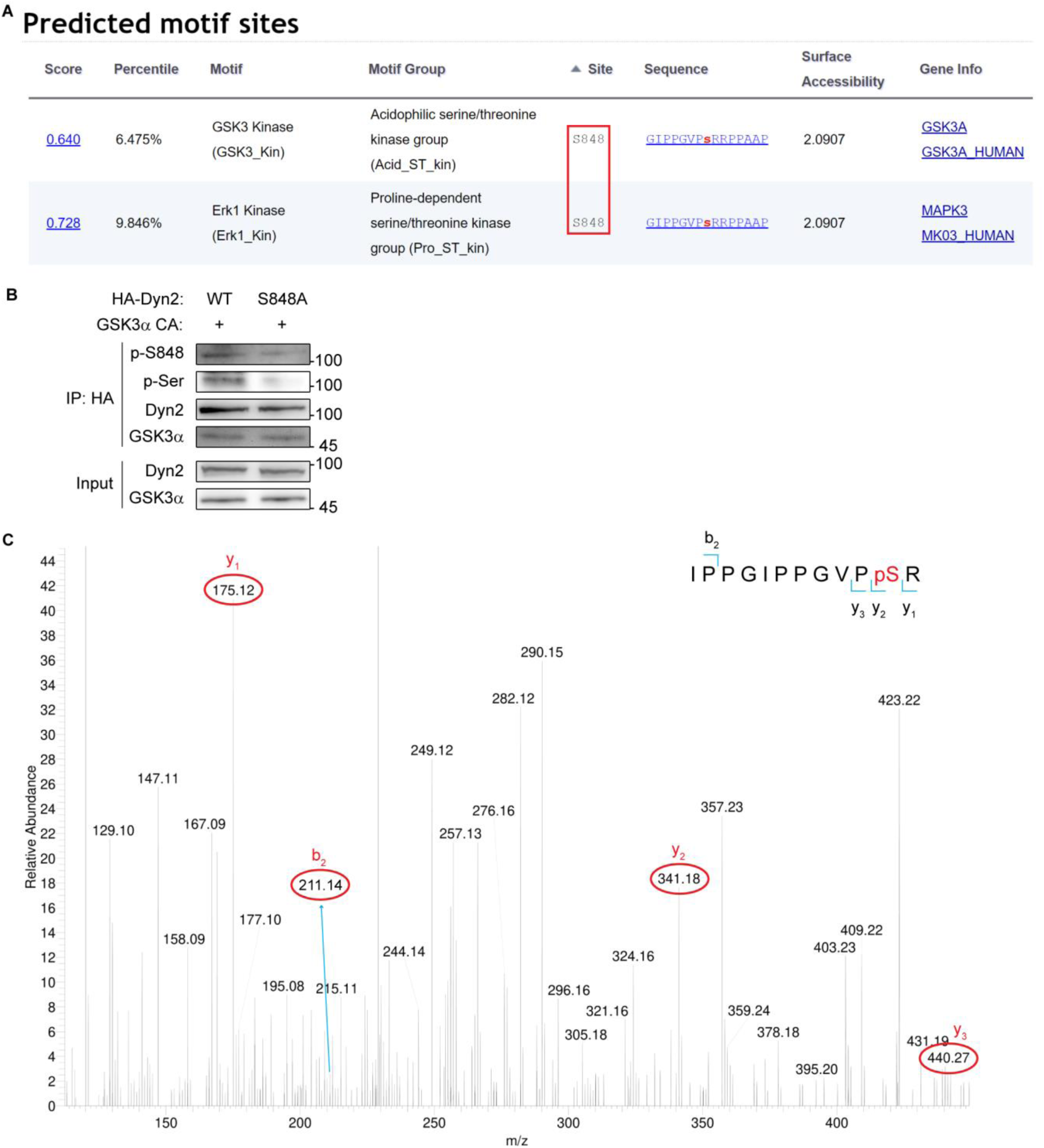
(related to Fig 6) GSK3α phosphorylates the S848 residue of Dyn2. (A) Analysis of potential kinases for Dyn2^S848^. Screenshot of the prediction results for Dyn2^S848^ kinases using scansite motif scans (https://scansite4.mit.edu/#scanProtein). Lower score indicates better match of protein sequence to the optimal binding pattern. (B) Analysis of Dyn2 phosphorylation under GSK3α overexpression. HA-Dyn2^WT^ or HA-Dyn2^S848A^ were co-expressed with GSK3α CA into Hela cells. After precipitated with anti-HA antibody, the phosphorylated Dyn2 was detected with Dyn2 phospho-S848-specific (p-S848) and anti-phospho-serine (p-Ser) antibodies. (C) Mass spectrometry analysis of Dyn2 phosphorylation by GSK3α. Recombinant Dyn2 was incubated with GSK3α and ATP in the *in vitro* kinase assay buffer for 1 hour. The reaction was then analyzed for phosphorylation with LC-MS/MS. The 838-IPPGIPPGVPpSR phosphopeptide containing phosphorylated Ser848 was identified. The MS/MS spectrum of this phosphopeptide is shown. The m/z of (3+) charged phosphopeptide is 422.89 with mass accuracy of <5 ppm. The mass difference between the y1 and y2 ions is consistent with phosphorylation at S848. Of note, the MS/MS spectrum is interfered by another peptide from Dyn2 (258-FFLSHPAYRH) with identical mass to our targeted phosphopeptide containing pS848.

## Supplementary videos

**Video S1. Time-Lapse Microscopy of Dyn2^WT^-mCherry and Bin1-GFP mutants in C2C12 myoblasts (related to Fig S1 I).** C2C12 myoblasts cotransfected with Dyn2^WT^-mCherry (magenta) and Bin-GFP mutants (green) were imaged with spinning disc confocal microscope with 5 s interval. The video shows the different dynamics between WT and CNM-Bin1 induced tubules under coexpression of Dyn2.

**Video S2. Time-Lapse Microscopy of Dyn2^WT^-mCherry mutants and Bin1^WT^-GFP in C2C12 myoblasts (related to Fig S2 D).** C2C12 myoblasts cotransfected with Dyn2^WT^-mCherry mutants (magenta) and Bin1^WT^-GFP (green) were imaged with lattice light sheet microscopy with 5 s interval.

## Supplementary tables

**Table S1.** List of plasmids used in this study.

| Plasmid | Vector | Source |
| --- | --- | --- |
| Dyn2 | pIEx-6 | Schmid lab<sup>#</sup> |
| Dyn2R465W | pIEx-6 | Chin et al., 2015 |
| Dyn2A618T | pIEx-6 | Chin et al., 2015 |
| Dyn2S619L | pIEx-6 | Chin et al., 2015 |
| Dyn2Δ625 | pIEx-6 | Chin et al., 2015 |
| Dyn2S848A | pIEx-6 | In this study |
| Dyn2S848E | pIEx-6 | In this study |
| Dyn2S856A | pIEx-6 | In this study |
| Dyn2S856E | pIEx-6 | In this study |
| Bin1 | pGEX-4T-1 | In this study |
| Bin1ΔSH3 | pGEX-4T-1 | In this study |
| Bin1ΔPI | pGEX-4T-1 | In this study |
| N-BAR | pGEX-4T-1 | In this study |
| Endophilin | pGEX | Schmid lab<sup>#</sup> |
| Bin1 | pET-30a | In this study |
| Bin1Q434X | pET-30a | In this study |
| Bin1K436X | pET-30a | In this study |
| Bin1-GFP | pEGFP-C1 | Addgene 22213 |
| Bin1Q434X-GFP | pEGFP-C1 | In this study |
| Bin1K436X-GFP | pEGFP-C1 | In this study |
| Dyn2-mCherry | pmCherry-N1 | Addgene 27689 |
| Dyn2S848A-mCherry | pmCherry-N1 | In this study |
| Dyn2S848E-mCherry | pmCherry-N1 | In this study |
| Dyn2G537C-mCherry | pmCherry-N1 | Chin et al., 2015 |
| HA-Dyn2 | pAS4.1w.Ppuro-aOn | In this study |
| HA-Dyn2S848A | pAS4.1w.Ppuro-aOn | In this study |
| HA-Dyn2S848E | pAS4.1w.Ppuro-aOn | In this study |
| GLUT4-HA-GFP | pQBI25 | McGraw lab* |
| HA-Dyn2 | pcDNA3.0-HA | In this study |
| HA-Dyn2S848A | pcDNA3.0-HA | In this study |
| GSK3A | pMT2 | Addgene 15896 |
| GSK3A-S21A | pMT2 | In this study |
| GSK3A-K148A | pMT2 | In this study |
| HA-GSK3B-S9A | pcDNA3 | Addgene 14754 |

#Dr. Sandra Schmid in the Department of Cell Biology, University of Texas Southwestern Medical Center, Dallas, TX 75390, USA
\*Dr. Timothy E. McGraw in Department of Biochemistry, Weill Cornell Medical College, New York, NY 10065

**Table S2.**
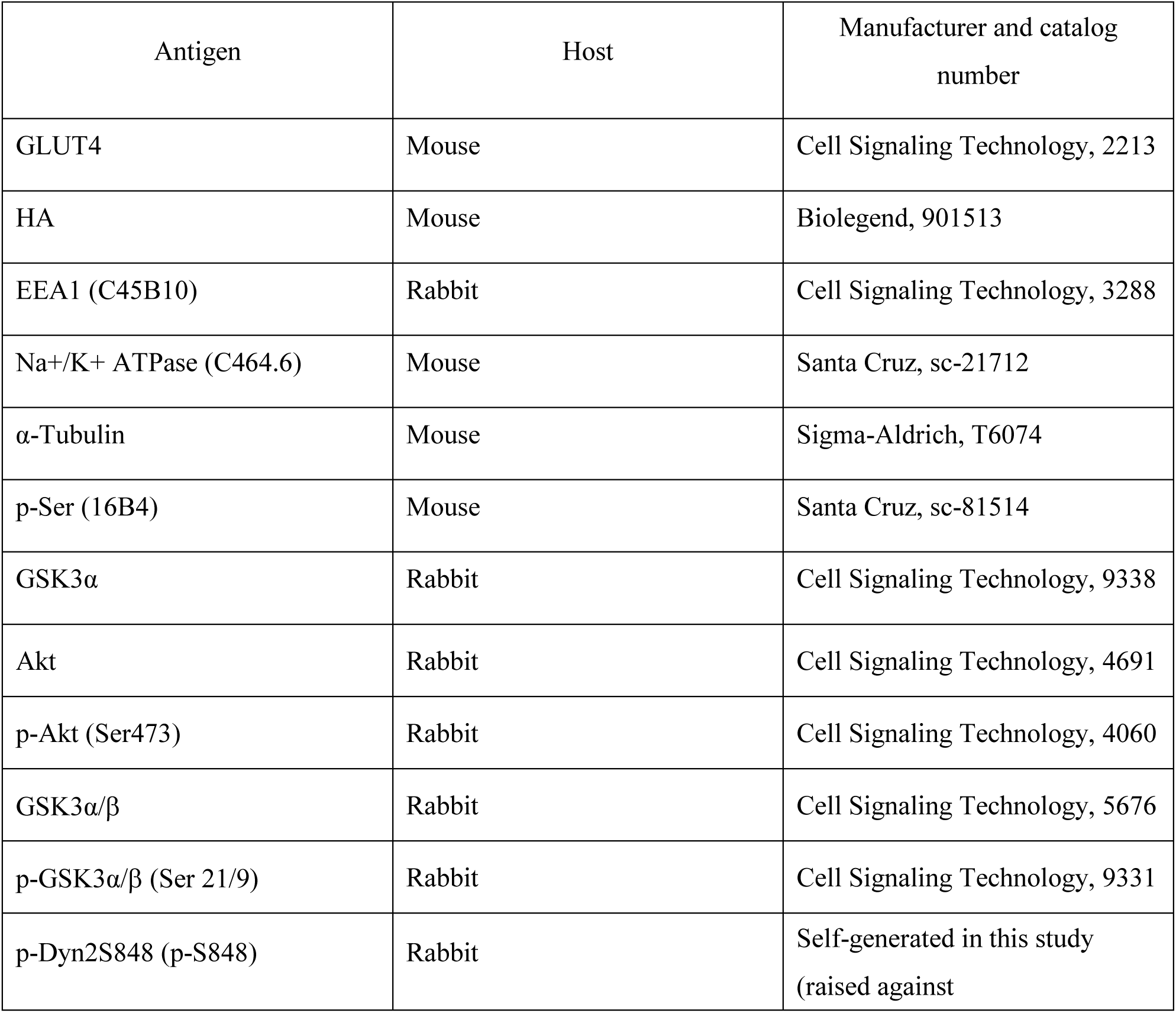
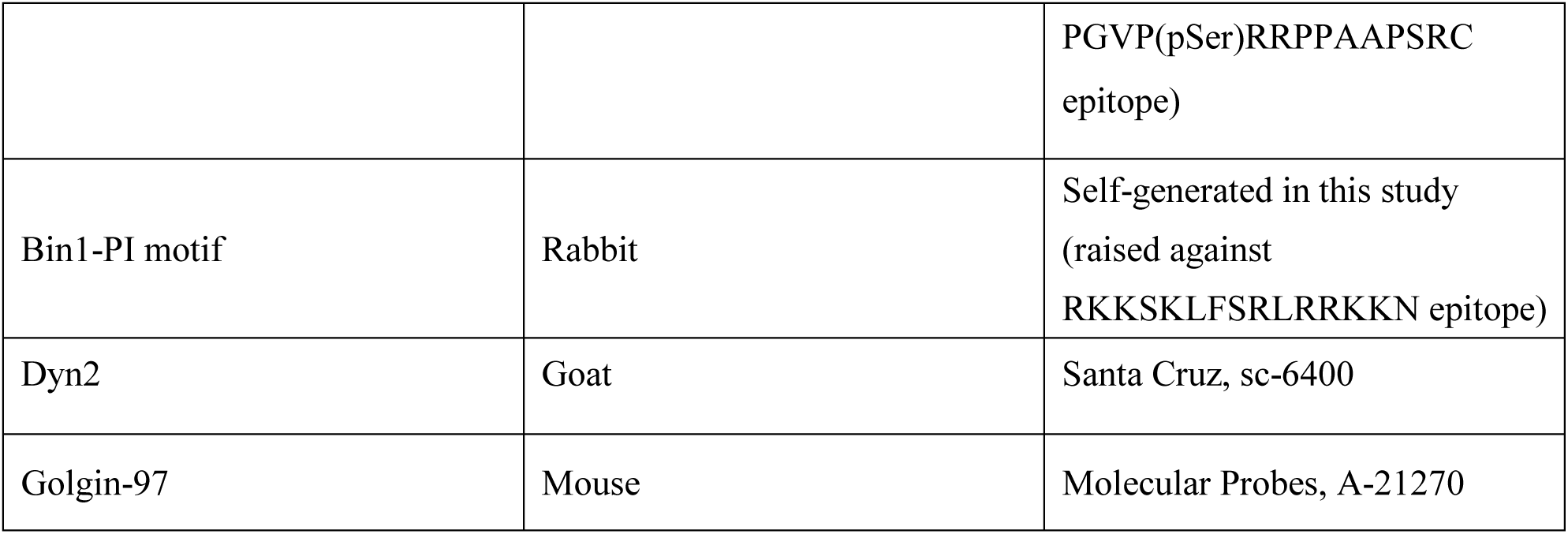
List of antibodies used in this study.

